# Calcium-binding protein expression alone is insufficient to identify and classify GABAergic neurons in macaque cortex

**DOI:** 10.64898/2026.01.26.701495

**Authors:** Alev M. Brigande, Juliane Krueger, Connor Park, Anita A. Disney

## Abstract

Understanding neuron subclasses and their functional consequences can contribute to understanding brain circuits. A scheme long used to classify GABAergic neurons in the neocortex is based on expression of three calcium-binding proteins (CBPs): parvalbumin (PV), calbindin D-28K (CB), and calretinin (CR). Because CB and CR are frequently co-expressed by individual neurons in rodents, this scheme has been replaced by one based on PV and two signaling peptides: somatostatin (SST) and vasoactive intestinal peptide (VIP). In macaques, however, CBPs are generally not co-expressed, and so their use has persisted despite suggestions that the underlying populations are not, in fact, entirely GABAergic. We set out to quantitatively evaluate CBPs as a classification scheme for GABAergic neurons in early and mid-level visual regions in macaque cortex. Combining immunohistochemistry and *in situ* hybridization, we find that up to half of neurons expressing CBPs are likely not GABAergic. Furthermore, contrary to what has been previously suggested, the GABAergic subpopulations cannot be distinguished based on staining intensity. Thus, the CBP-based classification scheme is not valid, at least as it has traditionally been used. Instead, we find support for co-labeling CB and CR neurons with SST and VIP, an approach that can identify GABAergic subpopulations within the CBP classes; or simply adopting the PV/SST/VIP scheme. We discuss the functional implications of expressing these various cell type markers, and how consideration of marker functions can support proper selection of a classification scheme for a given experimental purpose.

**Significance Statement:** The findings of this study challenge the currently dominant classification scheme for GABAergic neurons in macaque cortex, a scheme based on expression of the calcium-binding proteins parvalbumin, calbindin D-28K, and calretinin. Specifically, we show that a large proportion of neurons that express protein or mRNA for these calcium-binding proteins are likely not GABAergic. We go on to consider alternative approaches, discussing under what circumstances each might be useful.

## Introduction

A classification of neurons using anatomical criteria - such as morphology, protein/mRNA expression, or laminar position - can contribute to understanding circuit function, particularly when the classification system has functional relevance. One scheme that has been used for cortical GABAergic neurons is based on expression of three calcium-binding proteins (CBPs): parvalbumin (PV), calbindin D-28K (CB), and calretinin (CR; Kawaguchi and Kubota, 1993). This scheme has the advantage that - to the extent that we understand the consequences of expressing one CBP versus another (Schwaller et al., 2002; Schwaller, 2009) - it could offer an accompanying functional interpretation. However, for the resulting populations to constitute cell “classes”, marker expression should be mutually exclusive within individual neurons, at least at a given point in time. CBPs fail this test in rat cortex (Kawaguchi and Kubota, 1997), and so CBP-based classification was abandoned in rodents in favor of a scheme based on expression of one CBP (PV) and two signaling peptides: somatostatin (SST) and vasoactive intestinal peptide (VIP; Kawaguchi and Kubota, 1997). This PV/SST/VIP scheme has been a remarkably successful basis for describing inhibitory motifs in circuit models for rodent cortex (Pfeffer et al., 2013; Niell and Scanziani, 2021).

In macaques, however, evidence *against* co-expression of CBPs is strong for the primary visual cortex (V1; Kooijmans et al., 2020; Medalla et al., 2023) and probably for area 46 as well (Medalla et al., 2023). And so, with co-expression seeming species-specific, use of the PV/CB/CR scheme has continued in macaques (Van Brederode et al., 1990; DeFelipe, 1997; Meskenaite, 1997; DeFelipe et al., 1999; Kooijmans et al., 2020; Medalla et al., 2023). This despite reports that, in macaques, some CBP-expressing neurons may not be inhibitory (Van Brederode et al., 1990; Hof et al., 1999; Ichinohe et al., 2004; Constantinople et al., 2009). Recent studies (Kooijmans et al., 2020; Medalla et al., 2023) also disagree substantially with each other and prior literature regarding the relative proportions of CBP-immunoreactive (CBP-ir) subpopulations in V1 (Van Brederode et al., 1990; Meskenaite, 1997; DeFelipe et al., 1999; Kooijmans et al., 2020; Medalla et al., 2023). Furthermore, the composition of the CBP-expressing population appears to differ between V1 and prefrontal cortex (Van Brederode et al., 1990; Condé et al., 1994; Meskenaite, 1997; Medalla et al., 2023), leaving it unclear whether V1 is an oddball, or whether there may be some organizing principle; for example, an antero-posterior gradient in relative subpopulation sizes. Finally, beyond brief mention in a transcriptomic typing study (Chen et al., 2023), data on the relationship between CB-, CR-, SST-, and VIP-expressing neurons is lacking for macaques, limiting the ability to connect work in this critical translational species to work in rodents.

The extent to which CBP-based classification is justified for GABAergic neurons in macaque cortex, and what the resulting scheme can tell us about cortical circuits and their function, depends on a number of factors. First, is the scheme broadly valid: is a lack of CBP co-expression a feature of cortical neurons generally (i.e. beyond areas 17 [V1] and 46)? Then, if this is a valid scheme, one can ask *what is being classified*. Specifically, are most CBP-ir neurons GABAergic and, if not, can the GABAergic subpopulations be identified? Finally, one can ask whether CBP-based classification is useful: are the resulting classes functionally interpretable and/or correlated with other anatomically-defined classes?

To answer these questions, we quantify the extent to which PV-, CB-, and CR-expressing neurons in six early to mid-level visual areas of macaque cortex: 1) represent non-overlapping populations; 2) comprise similar proportions of the total CBP-expressing population; 3) are GABAergic (and whether inhibitory subpopulations can be identified); 4) account for the total GABAergic population; and 5) co-express SST and VIP. In doing so, we find that the manner in which this scheme has historically been used - to classify cortical GABAergic neurons in primates - is not justified.

## Methods

### Experimental design and statistical analysis

This manuscript describes two quantitative experiments. The first is a study of protein expression for PV-, CB-, and CR-expressing neurons in six visual cortical areas (V1, V2, V3, V3A, V4, MT/V5) in three macaques. We used triple immunofluorescence labeling with systematic random sampling of tissue, pseudo-random sampling of imaging sites, and quantification of all immunopositive neurons at a site (i.e. no sampling at the imaging site; n > 12,000 CBP-immunoreactive neurons from 26 tissue sections, 3 animals). The second is a study of mRNA expression in V1 and V2 of two macaques. We employed multiplexed *in situ* hybridization (RNAscope and MERFISH) for PV, CB, and CR mRNA along with markers for GABAergic neurons (glutamic acid decarboxylase type 1, GAD1, and the vesicular glutamate transporter, VGAT) and two signaling peptides (SST and VIP). Tissue sections for *in situ* were selected pseudo-randomly. The entire section (∼1.5 cm^2^) was imaged for MERFISH and all cortical cells counted (no sampling; n = 534,590 cells from 3 tissue sections, 2 animals). For RNAscope, imaging regions of interest were chosen pseudo-randomly and all cells within each ROI counted (n = 102,934 cells from 7 tissue sections, 1 animal). Analyses were performed on both per animal (n = animals) and pooled (n = neurons) data. Most measured quantities were not normally distributed, so tests are nonparametric unless otherwise specified.

### Animals

Tissue from six animals was used in this study. Two adult male (9 & 11 years; animal identifiers: A3 & A8) and one adult female (7 years; A9) *Macaca mulatta* and one juvenile male (3 years; A4) *Macaca nemestrina* were used in immunofluorescence experiments. These animals had all been used previously in unrelated *in vivo* recording experiments; for details of procedures in donor labs, see (Oristaglio et al., 2006; Lovejoy and Krauzlis, 2010; Nauhaus et al., 2012). Tissue from one experimentally naive adult female (11 years; A24) and one adult male (A25; also used in a V2-V1 tract-tracing study, opposite hemisphere) *Macaca mulatta* was used for *in situ* hybridization experiments. All procedures were performed in accordance with NIH guidelines for the humane care and use of animals, and approved by Animal Care and Use Committees at Columbia (A3, A8) and Duke (A24, A25) Universities and at the Salk Institute for Biological Studies (A4, A9).

### Fixed tissue harvest

Animals A3, A4, and A8 were transcardially perfused with heparinized 0.01M phosphate-buffered saline (PBS: pH 7.42, 0.9% NaCl, 1000 IU heparin/liter), followed by 4% paraformaldehyde (PFA) in 0.1M phosphate buffer (PB: pH 7.42). After removal from the skull, brain hemispheres were separated, blocked with ∼coronal cuts at the level of the lunate sulcus and anterior tip of the intraparietal sulcus, cryoprotected (immersion in 30% sucrose in PB), and stored at 4°C. Sections were cut at 50 μm on a freezing microtome and stored free-floating at 4°C, in PBS with 0.05% sodium azide added. Three 1-in-6 series of sections were set aside for each animal to make reference sets (Nissl, Gallyas, SMI-32) for defining laminar and brain area boundaries. Harvest for animal A9 was identical except 0.15% glutaraldehyde was included with the PFA in the fixative, and sections were therefore cut within 48 hours of perfusion and immediately reacted with 1% sodium borohydride in PB for 30 minutes, followed by 6 x 30-minute PB rinses.

### Fresh-frozen (unfixed) tissue harvest

Animals A24 and A25 were transcardially perfused with heparinized PBS (no fixation). After extraction, the brain was separated into hemispheres and the left (A24) or right (A25) hemisphere blocked (8-10 mm, ∼coronal plane) and immediately stored at -80°C. The other hemisphere was used in unrelated studies. The block containing the occipital pole was halved in the ∼horizontal plane and the ventral portion embedded in OCT embedding medium (Fisher). The tissue was cut at 10-12 μm, ∼coronal plane, using a cryostat. For four-plex fluorescence *in situ* hybridization (4-FISH, RNAscope; A24), sections were mounted on Superfrost Plus slides (VWR) and stored at -20°C for 1 hour, then at -80°C until use. For multiplexed fluorescence *in situ* hybridization (MERFISH, Vizgen; both animals), sections were mounted on MERSCOPE slides (Vizgen) and stored at -20°C for 5+ minutes before processing.

### Immunohistochemistry protocol optimization

We selected the dilution for each primary antibody based on dilution series experiments conducted in animal A3 using the ABC (avidin-biotin complex) method. Antibodies directed against PV, CB, and CR (see **Table 1**) were each tested at 1:125, 1:250, 1:500, 1:1000, and 1:2000 with all processing steps conducted at room temperature, and all incubations on a shaker. Sections were first blocked for 1 hour in 5% normal donkey serum (NDS; Jackson ImmunoResearch Labs, RRID:AB_2337258), 1% IgG-free bovine serum albumin (BSA; Jackson ImmunoResearch Labs, RRID:AB_2336947), 0.5% Triton X-100 (Sigma), and 0.05% sodium azide (Sigma) in PBS and then incubated for 24 hours in primary antibodies, at the various concentrations, diluted in blocking solution. After 3 x 10-15-minute PBS rinses, sections were incubated for 1 hour with the appropriate biotin-conjugated secondary antibody raised in donkey (see **Table 2**) diluted 1:1000 in PBS with 1% BSA added. After this incubation and rinsing (3 x 5 minutes), sections were incubated for 1 hour in an avidin-horseradish peroxidase solution (Vectastain Elite ABC kit, Vector Laboratories Cat# PK-6100, RRID:AB_2336819) and rinsed again in PBS (3 x 5 minutes). Antigenic sites were visualized using Vector’s VIP Substrate kit (Vector Laboratories Cat# SK-4600, RRID:AB_2336848), with reaction time controlled by eye (range: 3-11 minutes). After final PBS rinses, sections were mounted on gelatin-subbed slides and dried overnight. The next day, the tissue (now on-slide) was dehydrated through graded alcohols (50-100%) and 2 x 100% xylene, then coverslipped with Permount mounting medium (Electron Microscopy Sciences).

**Table 1:** Primary antibodies, all purchased from Swant.

| Target | Clonality | Host | Immunogen | Cat# | Lot # | Dilution | RRID |
| --- | --- | --- | --- | --- | --- | --- | --- |
| PV | poly | rabbit | rat muscle PV | PV25 | 5.10 | 1:1000 | AB_10000344 |
| CR | poly | goat | human recombinant CR | CG1 | 1§.1 | 1:250 | AB_10000342 |
| CB<br>(triple IF) | mono | mouse | chicken gut CB D-28k | 300 | 07(F) | 1:500 | AB_10000347 |
| CB<br>(double IF) | poly | rabbit | rat recombinant CB D-28k | CB-38a | 9.03 | 1:1000 | AB_3107026 |
| GABA | mono | mouse | GABA-BSA conjugate | 3A12 | ps2 | 1:5000 | AB_2314454 |

**Table 2:**
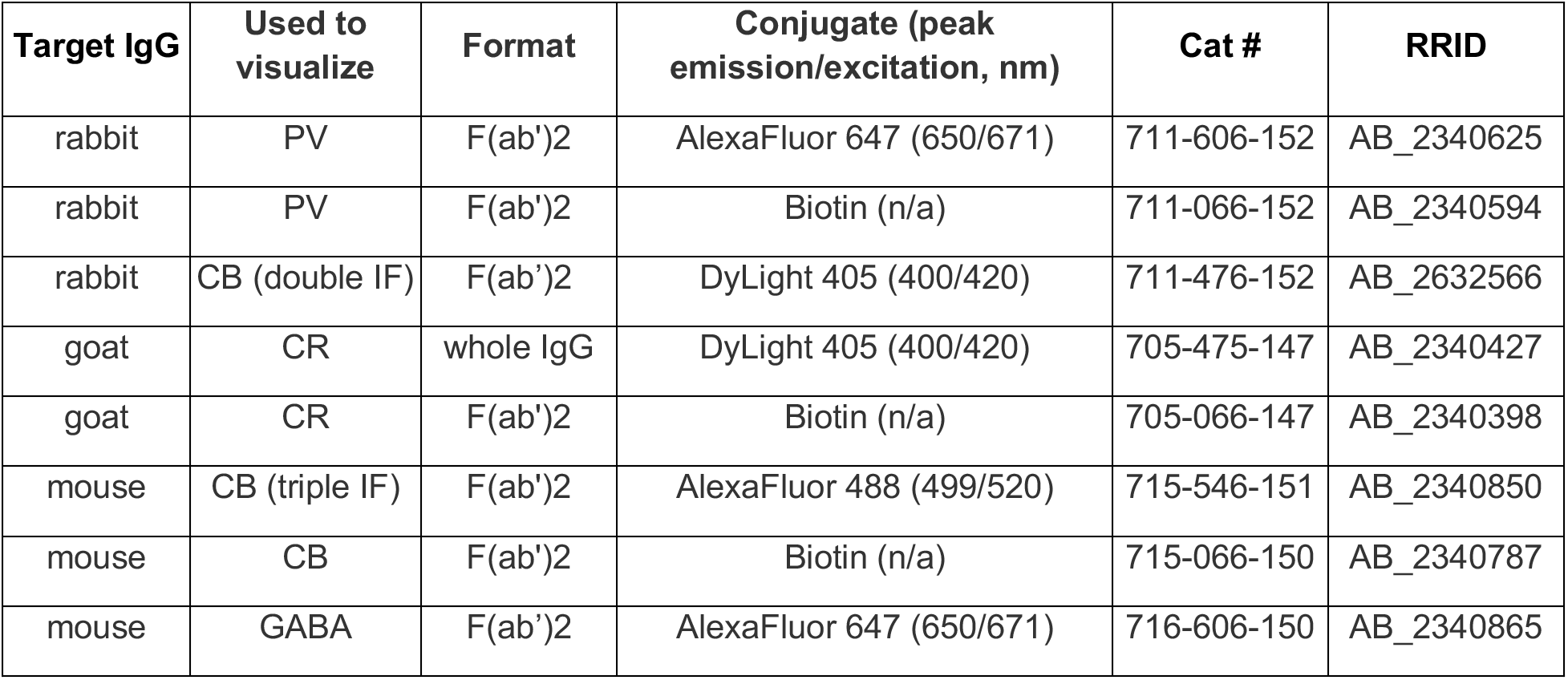
Secondary antibodies, all donkey host, Jackson ImmunoResearch.

| Target IgG | Used to visualize | Format | Conjugate (peak emission/excitation, nm) | Cat # | RRID |
| --- | --- | --- | --- | --- | --- |
| rabbit | PV | F(ab')<sub>2</sub> | AlexaFluor 647 (650/671) | 711-606-152 | AB_2340625 |
| rabbit | PV | F(ab')<sub>2</sub> | Biotin (n/a) | 711-066-152 | AB_2340594 |
| rabbit | CB (double IF) | F(ab')<sub>2</sub> | DyLight 405 (400/420) | 711-476-152 | AB_2632566 |
| goat | CR | whole IgG | DyLight 405 (400/420) | 705-475-147 | AB_2340427 |
| goat | CR | F(ab')<sub>2</sub> | Biotin (n/a) | 705-066-147 | AB_2340398 |
| mouse | CB (triple IF) | F(ab')<sub>2</sub> | AlexaFluor 488 (499/520) | 715-546-151 | AB_2340850 |
| mouse | CB | F(ab')<sub>2</sub> | Biotin (n/a) | 715-066-150 | AB_2340787 |
| mouse | GABA | F(ab')<sub>2</sub> | AlexaFluor 647 (650/671) | 716-606-150 | AB_2340865 |

The dilution that yielded the largest mean (non-stereological) count of cell bodies across ten, 250-μm-wide columns of tissue (for each antibody, each dilution) was tested in the triple immunofluorescence protocol described below, and a second non-stereological count made. For all three antibodies, mean counts (i.e. detection performance) did not differ between the best performing single-label immunohistochemistry condition and triple immunofluorescence (two-tailed t test, p > .05). This was the dilution used for the quantitative studies (**Table 1**).

### Antibody controls

The polyclonal antibodies used in quantitative triple IF experiments (rabbit anti-PV and goat anti-CR) both passed lot-specific preadsorption controls in macaque cortex.

### Triple immunofluorescence processing

A triple immunofluorescence (IF) protocol was employed against PV, CR and CB in animals A3, A4, and A8. A 1-in-24 series (1.2-mm section separation) of sections from the occipital pole to the anterior tip of the intraparietal sulcus was selected from a starting well that was randomized for each animal. All steps proceeded at room temperature and all incubations on a shaker. Sections were blocked for 1 hour in the solution described above (*Immunohistochemistry protocol optimization)* and incubated for 24-36 hours in a blocking solution containing all three primary antibodies diluted at the identified ‘optimal’ concentration (**Table 1**). Following this, sections were rinsed three times in PBS for at least 15 minutes each, before incubation for 6 hours in the dark with fluorophore-conjugated secondary antibodies raised in donkey (**Table 2**), diluted 1:400 in PBS with 1% BSA added. After this incubation and PBS rinses (3 x 10 minutes), sections were mounted on gelatin-subbed slides, dried overnight in the dark, dehydrated as above, and coverslipped with DPX mounting medium (Electron Microscopy Sciences). Coverslipped sections were stored at room temperature in the dark.

### Double immunofluorescence processing for GABA

A double IF protocol was employed against CB and GABA in animal A9. Two sections containing extrastriate cortex were pseudo-randomly selected and freeze-thawed (Disney et al., 2006). All steps proceeded at room temperature and all incubations on a shaker. Sections were blocked for 1 hour in 1% BSA in PBS with .05% sodium azide added, then incubated for 72 hours in blocking solution containing antibodies directed against CB and GABA (**Table 1**). Following this, sections were rinsed (3 x 15+ minutes) before co-incubation for 4 hours in the dark with fluorophore-conjugated secondary antibodies raised in donkey (**Table 2**), diluted at 1:200 (donkey anti-mouse-IgG) and 1:500 (donkey anti-rabbit IgG) in PBS with 1% BSA added. After this incubation and PBS rinses (3 x 10 minutes), sections were mounted on gelatin-subbed slides, dried overnight in the dark, dehydrated as above, and coverslipped with DPX mounting medium (Electron Microscopy Sciences). Coverslipped sections were stored at room temperature in the dark.

### Four-plex fluorescence in situ hybridization (4-FISH)

4-FISH (RNAscope, ACD) was performed on-slide in animal A24 using the RNAscope Multiplex Fluorescent Reagent Kit v2 and 4-plex Ancillary Kit. Custom probes targeted GAD1, VGAT, PV, CB, CR, SST, and VIP. See **Tables 3 and 4** for probe and fluorophore details and experimental design. The manufacturer’s protocol (Cat# UM323100, revision B, 2022) and 4-plex technical note (Cat# 323120-TN, revision A, 2019) were followed exactly with the following exceptions: sections were fixed in pre-chilled 4% PFA in PBS for 15 minutes at room temperature; the hydrophobic barrier air dried for 45 minutes; Protease III was used for 30 minutes at room temperature; slides were stored overnight in 5X saline sodium citrate at room temperature; Opal Dyes were diluted 1:1500; and ProLong Glass (Invitrogen) was used for mounting.

**Table 3:** 4-FISH probes, ACD.

| Gene | Target species | Channel(s) | Product name; Cat# | Lot # |
| --- | --- | --- | --- | --- |
| GAD1 | *M. nemestrina* | 3 | Mne-GAD1-C3; 1075571 | 21260D |
| VGAT | *M. mulatta* | 3 | Mmu-SLC32A1-C3; 1242311 | 23363C |
| PV | *M. mulatta* | 1, 4 | Mmu-PVALB-C1, C4; 461691 | 20050B, 23363C |
| CB | *M. mulatta* | 1, 2, 3 | Mmu-CALB1-C1, C2, C3; 461711 | 24002B, 23353B, 23363C |
| CR | *M. mulatta* | 4 | Mmu-CALB2-C4; 838461 | 23363C |
| SST | *M. mulatta* | 1 | Mmu-SST-C1; 461681 | 20211A |
| VIP | *M. mulatta* | 2 | Mmu-VIP-C2; 870571 | 20253D |
| *dapB* | bacteria | 1, 2, 3 | 3-plex Negative Control Probe; 320871 | 2017650 |

**Table 4:** 4-FISH experiment design.

| Slide number | Channel 1<br>(gene target, $\lambda$) | Channel 2<br>(gene target, $\lambda$) | Channel 3<br>(gene target, $\lambda$) | Channel 4<br>(gene target, $\lambda$) |
| --- | --- | --- | --- | --- |
| 7 | *dapB*, 520 | *dapB*, 570 | *dapB*, 650 | no probe, 620 |
| 9 | CB, 620 | VIP, 520 | GAD1, 570 | CR, 650 |
| 12 | CB, 620 | VIP, 570 | VGAT, 520 | CR, 650 |
| 16 | SST, 520 | VIP, 570 | CB, 620 | CR, 650 |
| 17 | SST, 620 | VIP, 650 | VGAT, 520 | PV, 570 |
| 18 | SST, 620 | VIP, 650 | GAD1, 570 | PV, 520 |
| 19 | PV, 520 | CB, 620 | GAD1, 570 | CR, 650 |
| 20 | PV, 570 | CB, 620 | VGAT, 520 | CR, 650 |

$\lambda$ : Opal fluorophore ~emission wavelength (Akoya). Peak excitation/emission ( $\lambda$ , nm): Opal 520, 494/525 (Cat# FP1487001KT, lot # 231221013); Opal 570, 550/570 (Cat# FP1488001KT, lot # 231213020); Opal 620, 588/616 (Cat# FP1495001KT, lot # 231016026); Opal 650, 627/650 (Cat# FP1496001KT, lot # 231023006).

### Multiplex in situ hybridization (MERFISH)

MERFISH (Vizgen) was performed on-slide in animals A24 (v1.0 chemistry) and A25 (v2.0 chemistry). A custom 300 Gene Panel (v1.0 chemistry: CM1560, Cat# 10400002; v2.0 chemistry: B2M2437, Cat# 10400179) was developed based on the *Macaca mulatta* genome and included probes for GAD1, VGAT, VGLUT1, CB, CR, SST, and VIP. A PV probe was requested in each panel. The v1.0 panel PV probe failed validation entirely (detection in nearly 100% of cells); the PV probe was re-made for v2.0 chemistry and was found to be selective, but insensitive. Panel 1 (A24) data for PV is not reported at all; Panel 2 (A25) data is not used for population quantification. Manufacturer protocols for tissue processing (v1.0 chemistry: Cat# 91600002; v2.0 chemistry: Cat# 91600132) and MERSCOPE Instrument use (Cat# 91600001) were followed exactly with the following exceptions: 4% PFA in PBS was pre-chilled (A24 only); photobleaching was performed for 5 (A24) or 12 hours (A25) in 70% ethanol; encoding probe hybridization was extended to 48-70 hours (A24 only); and larger coverslips were used (Bioptechs 22-mm round, VWR). Data were collected on the MERSCOPE Instrument with default parameters for a user-defined ∼1-1.5-cm^2^ region of interest (ROI) per slide that included V1 (both animals) and V2 (A24 only). Two ROIs (i.e., two separate processed slides) were imaged — at 7 z-axis planes per ROI — for A24, one for A25.

### Microscopy – IF

IF data were collected using Zeiss LSM 780 (triple IF) or 510 (double IF) confocal microscopes. Visual cortical areas were identified based on published criteria (Ungerleider and Desimone, 1986; Colby et al., 1988; Boussaoud et al., 1991). Within each brain area, imaging regions were selected pseudo-randomly. Laser lines (405, 488, 633 nm) were run at 2-20% power (level chosen to optimize signal-to-noise in the target channel while eliminating bleed-through to other channels). Other imaging parameters were held constant (40x water immersion objective, pinhole 49.5, pixel dwell 1.27 μsec, 8x line averaging). At each imaging location, two stacks of images at 1-µm intervals in the z-axis (’z-stack’) were made, one at just below the pia, the other just above the white matter. A single z-axis imaging plane was chosen from these z-stacks and a “tilescan” image captured of a ∼270-μm-wide region (or larger) that spanned all layers of cortex from pia to white matter. A minimum of three tilescans were collected per cortical area, per animal. In total, 54 tilescan ROIs were collected across 26 tissue sections. In all but one case (area V3 in A3, 2 sections), the three ROIs for a given brain area in a single animal came from three different tissue sections. ROIs for <u>different</u> brain areas often came from the same tissue section.

### Microscopy - 4-FISH

4-FISH data for V1 and V2 were collected using a Leica SP8 confocal microscope. Within each brain area, imaging regions were selected pseudo-randomly. The 405 laser and notched (488, 561, 594, 633 nm) white light sources were run sequentially between frames at 0.5-4% power, with power level chosen as above. Emission spectra were filtered between 415-480 nm for DAPI, 498-548 nm for Opal 520, 566-601 nm for Opal 570, 604-635 nm for Opal 620, and 643-673 nm for Opal 650. Other imaging parameters were held constant (20x oil immersion objective, pinhole 1 AU, pixel dwell 787 ns, 4x line averaging, frame size 2048 x 2048 pixels). Each of seven tissue sections was processed with a unique combination of mRNA probes (**Table 4**). For each of these seven experiments, two tilescan ROIs (single z-axis plane) were collected from V1, and one from V2. An additional section was processed with a negative control probe, from which one V2 tilescan was collected.

### Triple IF counting criteria and method

Manual counts and size measurements of immunoreactive (ir) somata were made using the measurement tool in Imaris (v10.2, Oxford Instruments) from isolated grayscale data channels (405/CR, 488/CB, 647/PV). Markers were placed to measure the long axis of the cell body; the number of these measurements provided the count. To be counted, a profile had to be filled with fluorophore (i.e. not punctate), in focus, and resemble a cell body in size and shape. All neurons were counted for comparisons between brain areas; neurons intersecting a 15-μm exclusion boundary around layer borders were excluded from laminar analyses. A randomly-selected (by animal, tilescan) population of 90 cell bodies for each population (CB-ir, CR-ir, PV-ir) were also measured perpendicular to their long axis, at the widest point (by eye), to yield aspect ratio data.

### Correction of 2D counts

In the Results below, we provide both raw population composition data, and estimates based on the Abercrombie correction:

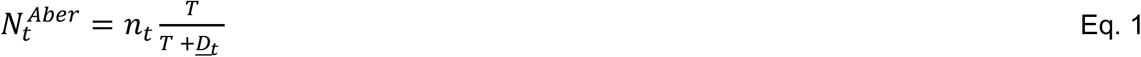

Where 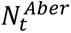 is the corrected count for cell type *t*, *n_t_* is the observed count, T is the tissue thickness, and *<u>D</u>_t_* is a measure of central tendency for cell diameter.

We observe ∼60% shrinkage in the z axis, in line with (Gardella et al., 2003). We used the above measurements of the long axis (*L_xy_*) of every counted soma to estimate original (pre-dehydration) soma diameters (*<u>D</u>_t_*) under isotropic x/y shrinkage ranging from 0 to 60%. A sensitivity analysis showed that results do not depend on x/y shrinkage (because *<u>D</u>_t_*<<T), so we set T = 50 μm (cut section thickness) for the Abercrombie correction. We used median (rather than mean) diameters (*<u>L</u>_xy_*; per cell type, by area) for *<u>D</u>_t_* in Eq. 1 because soma size distributions were non-normal (Shapiro-Wilk).

The Abercrombie correction assumes sphericity. To account for possible cell type-specific deviations from sphericity (e.g. bipolar neurons), somata were modeled as randomly oriented prolate spheroids with aspect ratio 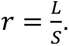. For a spheroid with long-axis diameter *L*_0_ = *D*_0_ and short-axis diameter 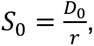, the z-direction intercept for a given orientation vector is:

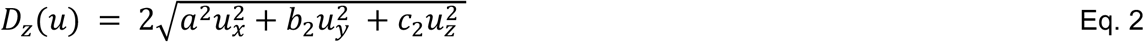

with semi-axes 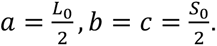. Assuming uniformly random orientations, the expected z-axis profile of a neuron is then:

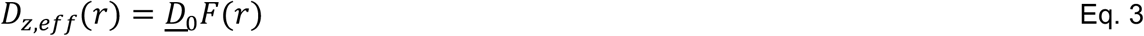

where

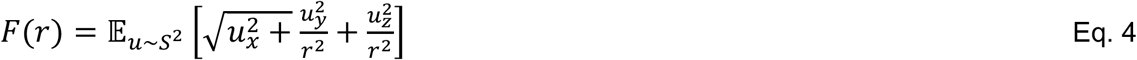

is a dimensionless factor computed once per aspect ratio using Monte Carlo simulation (200,000 random orientations). For *r* = 1 (i.e. spheres), *F*(*r*) = 1; for elongated somata, *F*(*r*)< 1. Ellipsoid corrected Abercrombie counts were computed by substituting *D_z_*_,*eff*_(*r*) for *<u>D</u>_t_* in Eq.1.

### Bimodality analysis of triple IF intensity

Fluorescence intensities were quantified for individual neurons using median pixel intensity values within a manually-centered somatic region of interest (ROI) for every counted neuron. We first assessed evidence for bimodality in individual tilescans. For each image and cell type with n > 50 neurons, we fit one- and two-component Gaussian Mixture Models (GMMs) to the raw intensity data and compared models using a Bayesian Information Criterion (BIC) difference measure (ΔBIC = BIC1 - BIC2, values > 0 support a two-component model), supplemented by Ashman’s D as a measure of component separation. The median values for ΔBIC (12.6) and Ashman’s D (2.42; 75th percentile, 3.56) supported bimodality, so we continued with population analysis.

To remove image-to-image variation, median intensities were log-transformed, z-scored, and pooled across animals and brain areas. For each cell type, we fit one- and two-component GMMs to the pooled data, quantified ΔBIC, and computed Ashman’s D and Hartigan’s dip statistic. Finally, if a 2-component model was supported, we estimated the proportion of high-intensity neurons using thresholds derived from receiver operating characteristic (ROC) analyses performed on the normalized intensity distributions.

### 4-FISH analysis

Automated counting with manual curation was performed in Imaris. DAPI-stained nuclei were segmented using Cellpose-SAM (v4.1.1; Pachitariu et al., 2025) with default parameters, imported as Surfaces, and filtered (number of voxels ≥ 310, equivalent to area ≥ 25 um^2^). Fluorescent puncta (mRNA transcripts) were detected using the Spots tool with background subtraction and an estimated diameter of 1 μm. For each channel, puncta were filtered using the “quality” metric to include threshold cases (minimum quality, 3.5-20). Puncta in the 520 and 620 channels were then classified using ML to exclude lipofuscin. Nuclei and puncta were imported into the Cells tool to determine ∼puncta/nucleus, and the counts were then exported along with x/y position. Laminar borders were drawn as connected segments using Measurement Points, and x/y positions exported. Cells were assigned to cortical layers, and the approximate area of each layer computed. All cells not in white matter were counted for comparisons between brain areas; cells intersecting a 15-μm exclusion boundary around layer borders were excluded from laminar analyses.

### MERFISH analysis

Automated image processing was performed using MERSCOPE software (v234b.241217.1593, Vizgen) with decoding algorithm 1-1 and segmentation algorithm Cellpose (Stringer et al., 2021). Segmented cells passed quality control (QC) if they met three criteria: 1) volume: 100-3000 um^3^ (Zhang et al., 2023), 2) 5+ gene transcripts detected (more permissive than Yao et al., 2023), and 3) blanks < 10% of detected transcripts (false positives). Cells were assigned to V1 or V2 and to cortical layers based on hand-drawn ROIs using MERSCOPE Vizualizer (v2.5.3503.0, Vizgen). All cells were counted for comparisons between brain areas and for laminar analyses.

### Binarization of ISH counts

To classify a cell as positive or negative for a gene, we set a copy-number threshold separately for each gene within each detection method (i.e., one threshold per gene across all slides for 4-FISH and one threshold per gene for each of the two MERFISH chemistries). We first estimated the background distribution of spurious counts by fitting a negative binomial model (or Poisson model, where counts were not over-dispersed) to the low-count cells, defined by an elbow rule as those with counts below the smallest count k ≥ 2 whose relative frequency is at least half that of count k-1 (the point at which the histogram’s decay flattens); if no such k exists, k = 2. We then set the binarization cutoff at the smallest copy number for which the expected background false positive rate (FPR) fell below 1%. This approach ensures that calling a cell “positive” carries the same statistical meaning, regardless of probe or chemistry, and thus allows us to combine data across experiments to increase power.

### Bimodality analysis of MERFISH copy counts

We repeated the mixture-model analysis on MERFISH copy-count data from animals A24 and A25. Cells were considered CB- or CR-positive if they contained ≥ 2 copies of the respective transcript (a flat cutoff). For each marker we fit one- and two-component GMMs to the copy-count data and compared models by ΔBIC, supplemented by Ashman’s D, as above; unlike the intensity data, counts were modeled directly, without log-transformation or z-scoring, as copy number is already comparable across images and animals.

GAD1 and VGAT were used as a molecular ground truth for GABAergic identity, with cells called GAD1+ or VGAT+ at the per-animal copy-number thresholds defined above (Binarization of ISH counts). Taking the posterior probability of the high-count GMM component as a classifier score, we computed ROC curves and areas under the curve (AUC) against three definitions of “GABAergic” — GAD1+, VGAT+, and both GAD1+ and VGAT+ — and applied a cost-sensitive Bayes rule, assigning a cell to the high-count class when the posterior P(high | x) was at least a threshold τ = C(FP)/[C(FP) + C(FN)].

### Image adjustments in producing Figures

In producing figures for publication, recoloring and contrast and brightness adjustment were applied to improve perceptibility for color anomalous readers. Note that these steps were taken for publication only; <u>all</u> data were collected from single channel, grayscale images, unadjusted.

### Software and code

Analyses were conducted in Python 3.11/3.12 and in R 4.5.2.

## Results

All neurons immunoreactive (-ir) for PV, CB, and CR (i.e. PV-ir, CB-ir, CR-ir neurons) in tilescans of V1, V2, V3, V3A, V4, and MT/V5 (**Figure 1**; **Figures S1, S2, S3**) were measured along the long axis of their soma, and counted. Pooling the raw counts across all six brain areas (152.73 mm^2^): of 12,247 neurons, 6,443 (52.6%) were PV-ir, 3,460 (28.3%) were CB-ir, and 2,344 (19.1%) were CR-ir. Abercrombie-corrected percentages and long axis measurements for the full population, and aspect ratios for a sample of 270 neurons, are shown in **Table 5**. Abercrombie-corrected counts yield nearly identical percentages both under assumptions of sphericity (PV-ir = 51.9%, CB-ir = 28.4%, CR-ir = 19.7%) and for a ‘worst case scenario’ ellipsoid model (PV-ir = 51.6%, CB-ir = 28.6%, CR-ir = 19.8%) in which PV-ir neurons are modeled at their average aspect ratio (1.7), and CB- and CR-ir neurons are modeled at their respective maximum measured aspect ratios (2.4 and 3.3).

**Figure 1:**
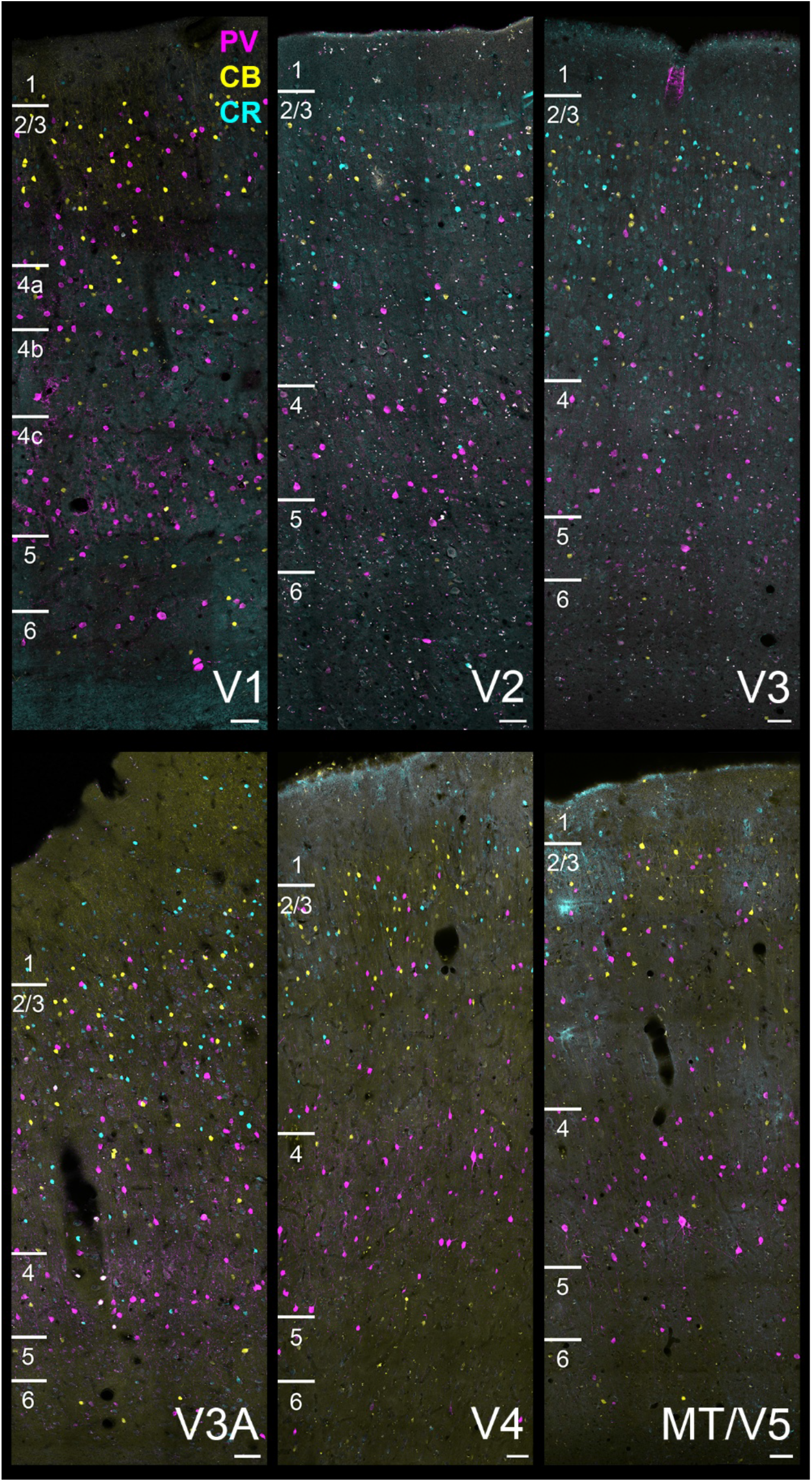
Triple immunofluorescence labeling for CBPs in early to mid-level visual cortex. Immunofluorescence is shown for PV (magenta), CB (yellow) and CR (cyan) in cortical areas V1 (top left) through MT/V5 (bottom right). Cortical layer borders are indicated to the left of each panel; note that layer boundaries in the tissue are not parallel with image borders/layer markers in all panels. Tissue shown covers all three animals. A3: V4, V5; A4: V1, V3A; A8: V2, V3. Scale bar: 50 μm, all panels.

**Table 5:**
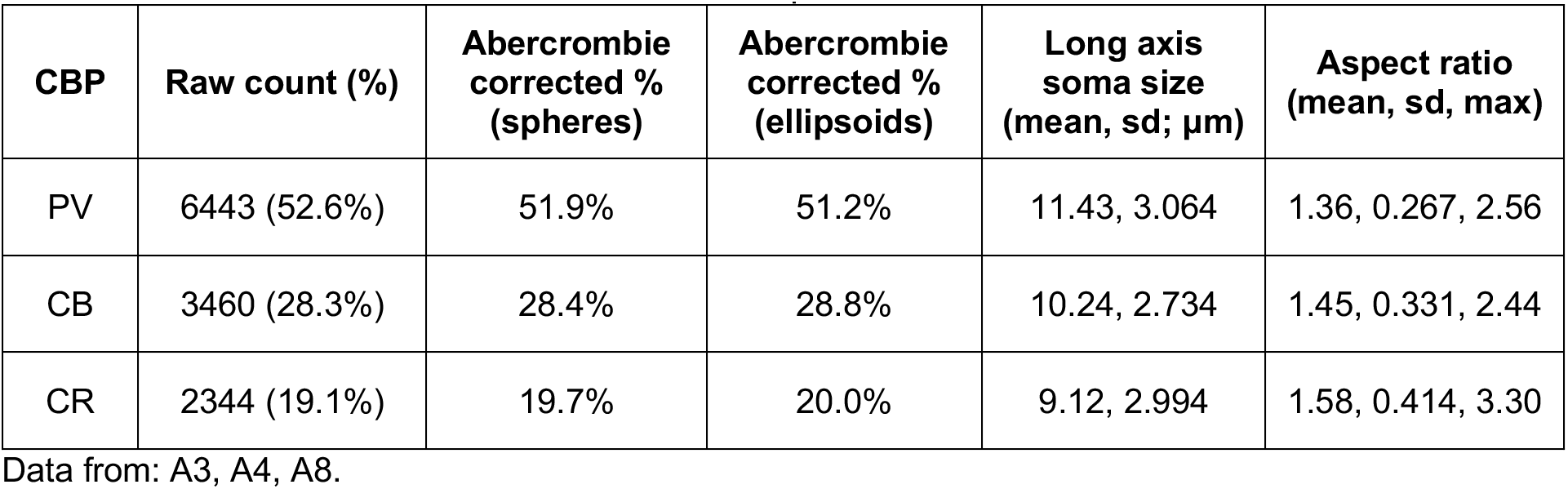
Pooled count data and statistics for Triple-IF detection of PV, CB, and CR.

Subtype composition differed across visual areas (**Figure 2A**). To assess area differences in these compositional data, we transformed area-level pooled subtype proportions using the centered log-ratio transform, and computed pairwise Aitchison distances (**Figure 2B**; Pawlowsky-Glahn and Egozcue, 2006; Quinn et al., 2019). Low Aitchison distances (dark cells in the matrix in **Figure 2B**) indicate greater compositional similarity. Under this analysis, V2 and V3 are most similar; then MT/V5 clusters more closely with these two early visual areas than with V3A or V4, which are dissimilar to each other. V1 is compositionally distinct from all extrastriate areas.

**Figure 2:**
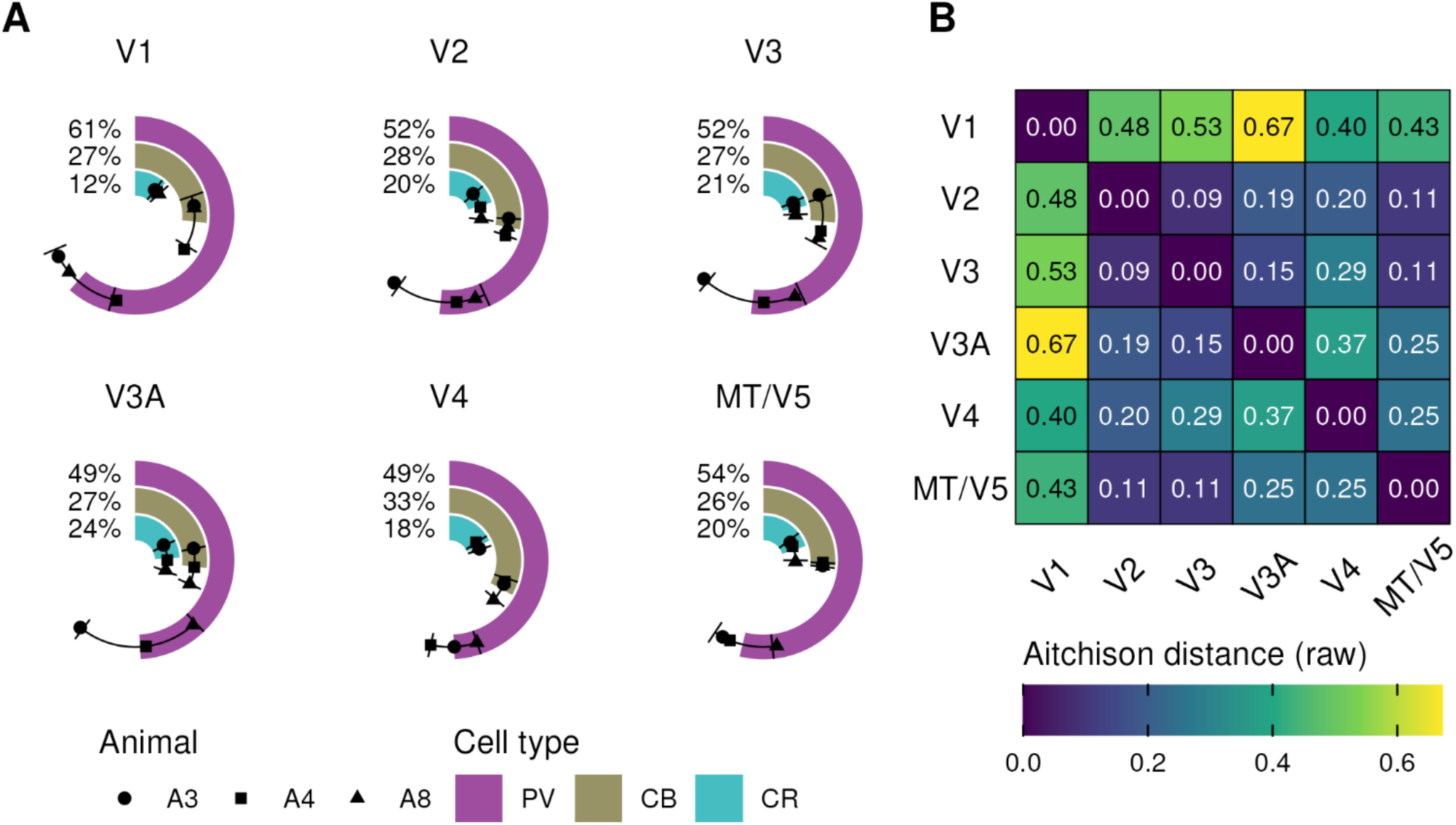
Composition of CBP-ir populations across early to mid-level visual cortex. (A) Radial bar graphs (in which a bar that makes full circle would indicate that a given cell type accounts for 100% of neurons in a brain area; bars sum to 100) of the population composition (mean, sd) for each visual cortical area show that CB-ir neurons make up ∼one quarter of CBP-ir neurons in all areas except V4, where they account for one third. PV-ir neurons comprise ∼half or more of CBP-ir neurons in all areas, and a particularly large proportion in V1. CR-ir neurons are particularly sparse in V1, and account for one fifth to one quarter of CBP-ir neurons in extrastriate cortex. (B) These patterns are reflected in the Aitchison distances between areas, which show that V1 (yellow/green shades) is distinct from all other visual cortical areas. Data from: A3, A4, A8. Area sampled, number of neurons: V1 = 6.243 mm^2^, 1,383 neurons; V2 = 19.293 mm^2^, 1,769 neurons; V3 = 11.328 mm^2^, 1,741 neurons; V3A = 12.906 mm^2^, 2,041 neurons; V4 = 97.447 mm^2^, 2,927 neurons; MT/V5 = 5.512mm^2^, 2,386 neurons.

Interestingly, CB-ir neurons comprise a strikingly similar proportion (26-28%) of CBP-ir neurons across areas, with the possible exception of area V4 (33%). In each animal individually, the proportion of CB-ir neurons was higher (1.5-8%) in V4 than the average across other visual areas. Given the small number of animals, this effect did not reach statistical significance in an exploratory paired permutation test (p ≈ 0.25). Biased sampling of layers 2 and 3 could masquerade as compositional differences between brain areas, due to differences in laminar composition (see below); most CB-ir (and CR-ir) neurons are found in these superficial layers. For area V4, layers 2 and 3 comprise a slightly <u>smaller</u> proportion of the sampled tissue (35.8%, 1.5 sd from the mean across areas) than they do for other areas (39.2-44.3%), which would actually tend to favor <u>lower</u> counts for CB-ir neurons.

Published data on the composition of the CBP-ir population in V1 have differed substantially from each other, and in some cases from our data. As an independent measure, we compiled equivalent data from our 4-FISH study of mRNA expression (**Figure 3, top; Figure S4, top**) in a different animal (A24). We find similar - but not identical - proportions in V1 to those we observed for protein expression, with a notably large CR-ir population (**Figure 3, top; Figure S4, top**): of 2,177 V1 neurons expressing mRNA for a CBP (CBP+ neurons), 56% express PV mRNA (PV+; n = 1,224), 30% express CB mRNA (CB+; n = 660), and 23% express CR mRNA (CR+; n = 507). In the text that follows, CBP-ir (and PV-ir, etc.) refers to detection of protein (IF) and CBP+ (PV+, etc.) to detection of mRNA.

**Figure 3:**
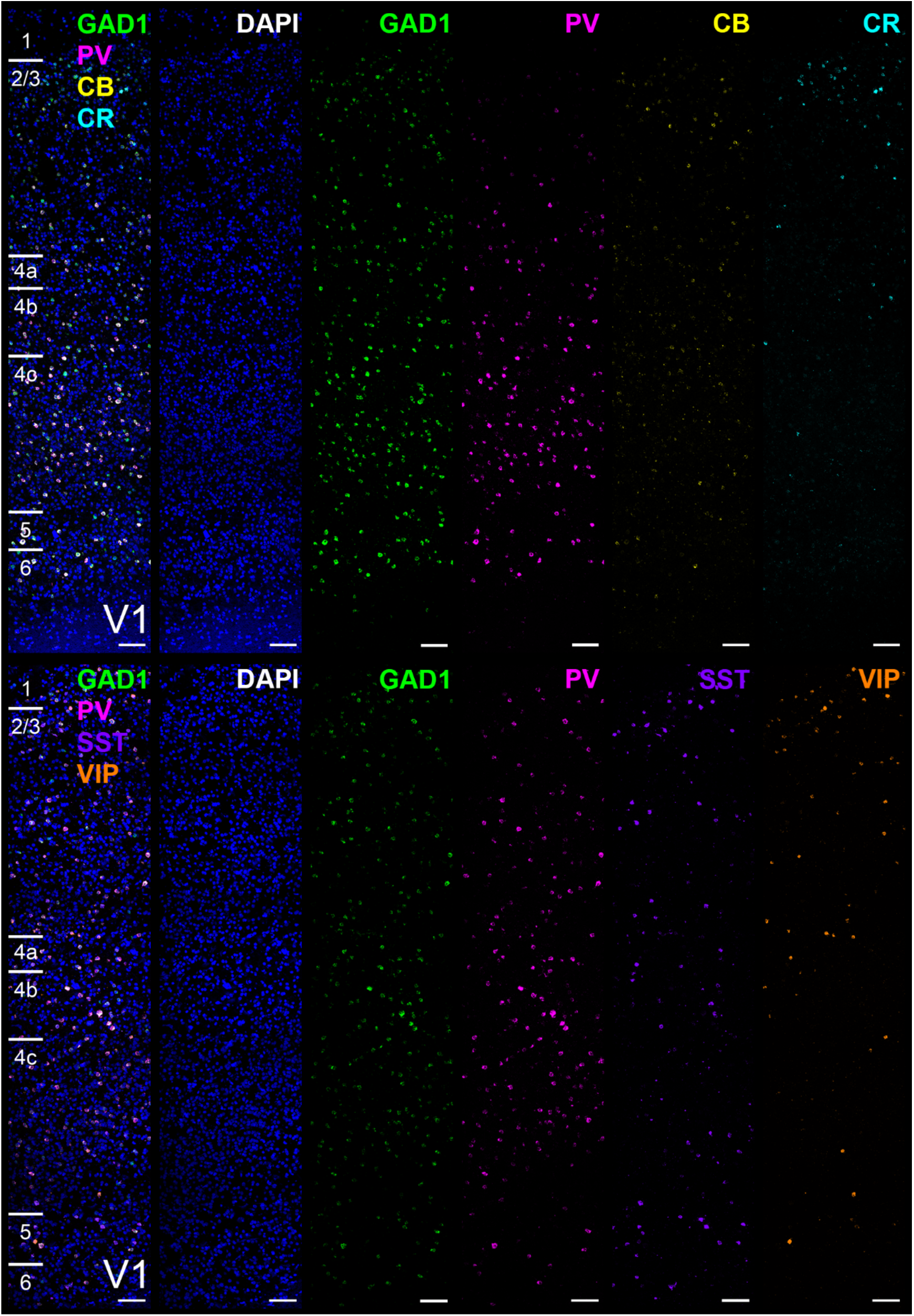
4-plex RNAscope fluorescent labeling (4-FISH) for CBP (top row) and neuropeptide (bottom row) mRNA transcripts in cortical area V1. DAPI-stained nuclei (blue) with fluorescent puncta (mRNA transcripts) are shown for GAD1 (green), PV (magenta), CB (yellow), CR (cyan), SST (purple), and VIP (orange). Cortical layer borders are indicated to the left of each panel; note that layer boundaries in the tissue are not parallel with image borders/layer markers in all panels. Experimental design in **Table 4**. Data from: A24. Scale bar: 100 μm, all panels.

The composition of the CBP-ir (i.e. protein expression) population also differs by cortical layer (**Figure 4A**), and the laminar pattern does not differ between brain areas (Kruskal-Wallis, minimum value p = 0.0856, maximum p = 0.8684; heatmap of proportions, **Figure 4B**). In this statistical comparison, V1 layers 4a and 4b (**Figure 4C**) were left out of the analysis and only layer 4c was compared against extrastriate layer 4. This analysis excluded 518 neurons (across brain areas, ∼4%) that were within a 15-μm confidence boundary around each layer border.

**Figure 4:**
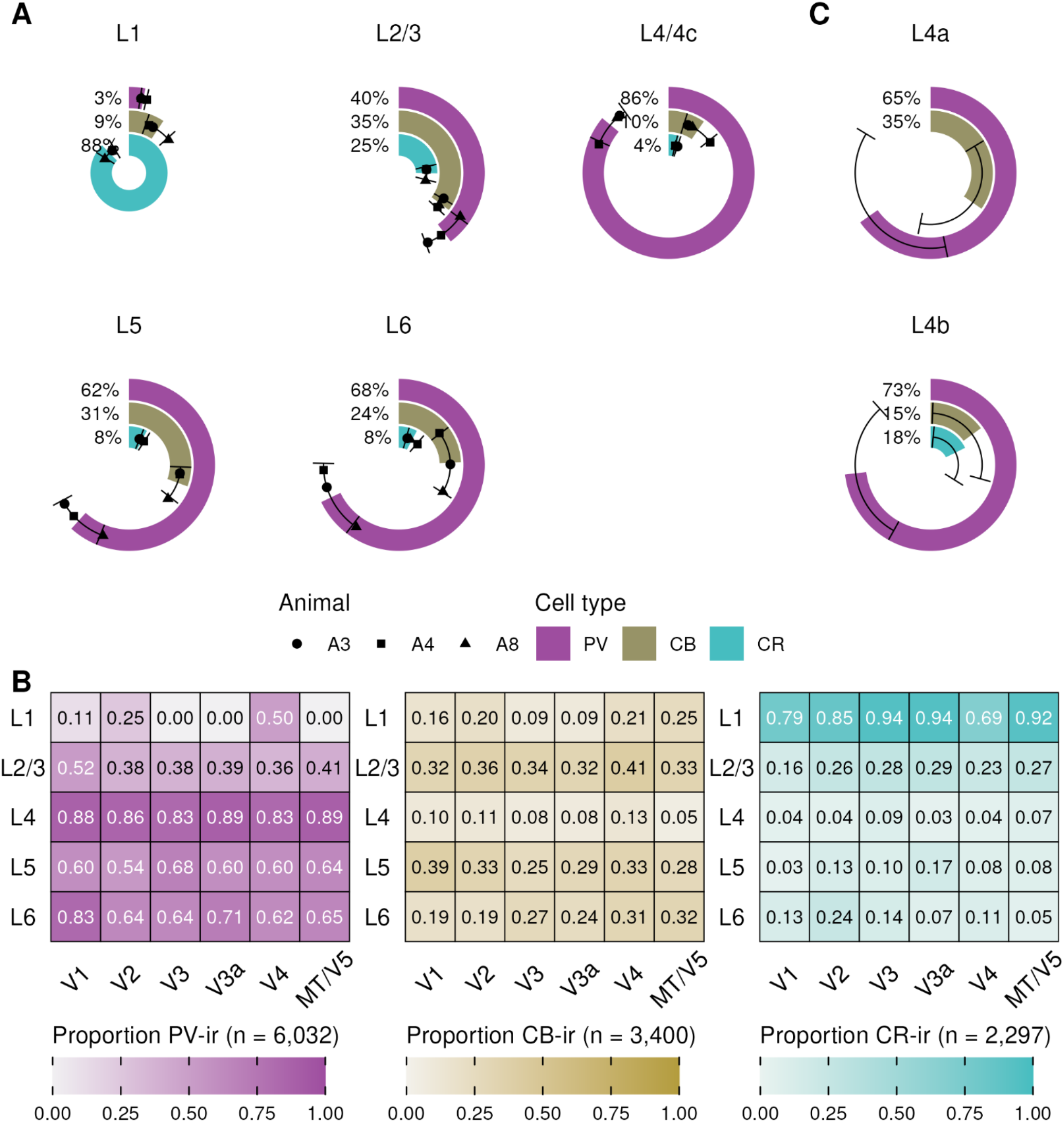
Composition of CBP-ir population by layer across early to mid-level visual cortex. (A) Radial bar plots for each layer (mean, sd; combined across brain areas) show that CR-ir neurons dominate in layer 1, and PV-ir neurons in layers 4/4c to 6. Layers 2 and 3 have a roughly balanced population of all three cell types. (B) Count heatmaps show that this laminar motif is similar across brain areas. In this panel, for V1, ‘L4’ is layer 4c only. (C) Radial bar plots for layers 4a and 4b of V1, which strictly have no anatomical equivalent in extrastriate areas, show they are more like layer 4c than layers 2 and 3. Data from: A3, A4, A8.

For neurons defined by CBP immunoreactivity to constitute inhibitory cell classes (as they have generally been used in anatomical studies of macaque cortex), they should represent i) largely non-overlapping populations of ii) GABAergic neurons. It also seems desirable that such a scheme iii) account for a substantial fraction of the total GABAergic population, and perhaps also iv) be functionally interpretable. We address each of these four criteria below.

### Criterion 1: Neurons immunoreactive for parvalbumin, calbindin, and calretinin comprise largely non-overlapping populations in early and mid-level visual cortical areas

Across all areas combined, only 485 (4.0%) of 12,247 CBP-ir neurons across animals A3, A4, and A8 were immunoreactive for two or more CBPs. 271 (2.2%) expressed both PV and CB, 154 (1.3%) expressed PV and CR, and 27 (0.2%) expressed CB and CR. 33 neurons (0.3%) expressed all three CBPs. All types of double- and triple-labeled cells were seen roughly equally often in all brain areas. PV/CR and CB/CR types were most often encountered in layers 2 and 3 (65% of PV/CR cells; 85% of CB/CR cells). In MT/V5 and V3A, PV/CB cells were mostly found in layers 2,3, and 5 (78% for MT/V5; 71% for V3A). In other areas, PV/CB cells were roughly evenly distributed across layers, as were PV/CB/CR neurons in all areas.

We also assayed CBP mRNA co-expression in V1 and V2 in A24 by 4-FISH and found similarly low levels (**Figure 5**; direct comparison to Triple-IF: **Figure S5**; count data: **Table S1**). Across V1 and V2 combined, of 3,409 CBP+ neurons, 316 (9.3%) were positive for two or more CBPs: 133 expressed PV and CB (3.9%; compared to 2.2% for protein), 126 expressed PV and CR (3.7%; 1.3% for protein), and 45 expressed CB and CR (1.3%; 0.2% for protein). Twelve neurons were positive for all three CBPs (0.4%; 0.3% for protein).

**Figure 5:**
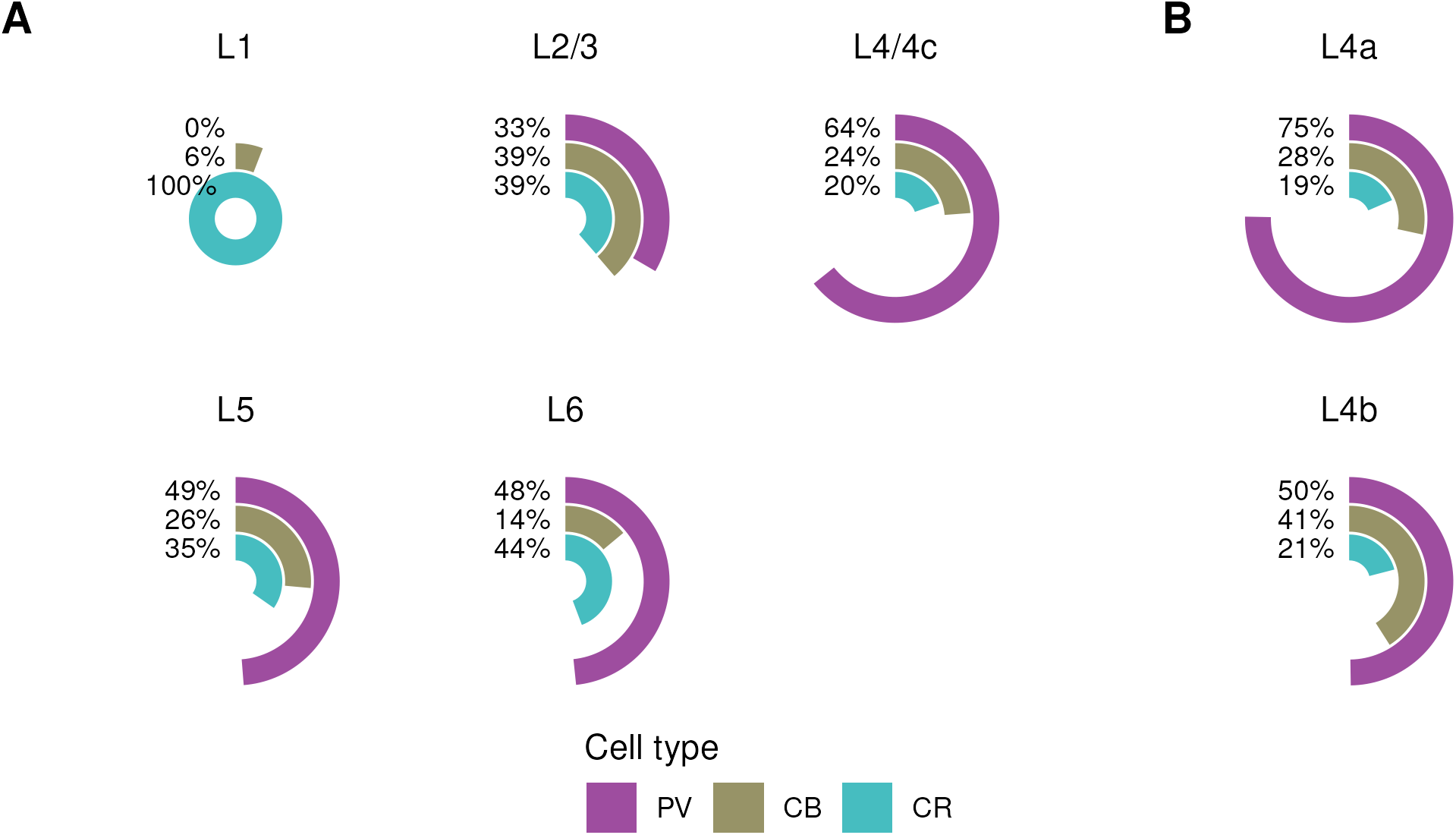
Composition of CBP+ population by layer in V1 and V2, by 4-FISH detection of mRNA. (A) Radial bar plots for each layer show that, as for protein expression shown in Figure 4, CR+ neurons dominate in layer 1, and PV+ neurons are the larger populations in layers 4/4c to 6. Layers 2 and 3 have a roughly balanced population of all three cell types. (B) Radial bar plots for layers 4a and 4b of V1 show that when mRNA is measured, layer 4a is similar to layer 4c, and 4b is similar to layers 2 and 3. Data from: A24.

### Criterion 2: A substantial fraction of neurons immunoreactive for parvalbumin, calbindin, and calretinin are <u>not</u> GABAergic

It has long been known that the CBP-based classification scheme, at least in a strict interpretation, fails this criterion; PV (Van Brederode et al., 1990; DeFelipe, 1997; Ichinohe et al., 2004; Constantinople et al., 2009), CB (Van Brederode et al., 1990; DeFelipe, 1997), and CR (DeFelipe, 1997; Meskenaite, 1997) are all expressed by excitatory neurons in the cortex of macaques. Past studies that relied on immunolabeling (protein expression) to identify GABAergic neurons used antibodies directed against different targets: either the synthetic enzyme GAD (glutamic acid decarboxylase) or against GABA itself. When colchicine is not used pre-mortem to stop axonal transport, antibodies directed against GAD have a high rate of false negatives for soma counting (∼80%, our own unpublished data). Antibodies directed against GABA, on the other hand, call for the use of glutaraldehyde as a fixative, which leads to false negatives for detection of CBPs (Meskenaite, 1997; Disney and Reynolds, 2014).

To address these limitations, we used 4-FISH detection of mRNA, instead of traditional protein immunolabeling, in animal A24 and found that putatively excitatory neurons (GAD1 or VGAT mRNA-negative) make up a substantial proportion of CBP+ cells in both V1 and V2 (**Figure 6**; **Tables S2, S3**). In separate highly multiplexed (MERFISH) mRNA detection experiments, just under half of the putatively inhibitory (GAD1+ and/or VGAT+) neurons expressed the type 1 vesicular glutamate transporter (VGLUT1; **Table 6**). These neurons - which may have dual release capabilities - account for ∼10% of the VGLUT1+ population (**Table S4**).

**Figure 6:**
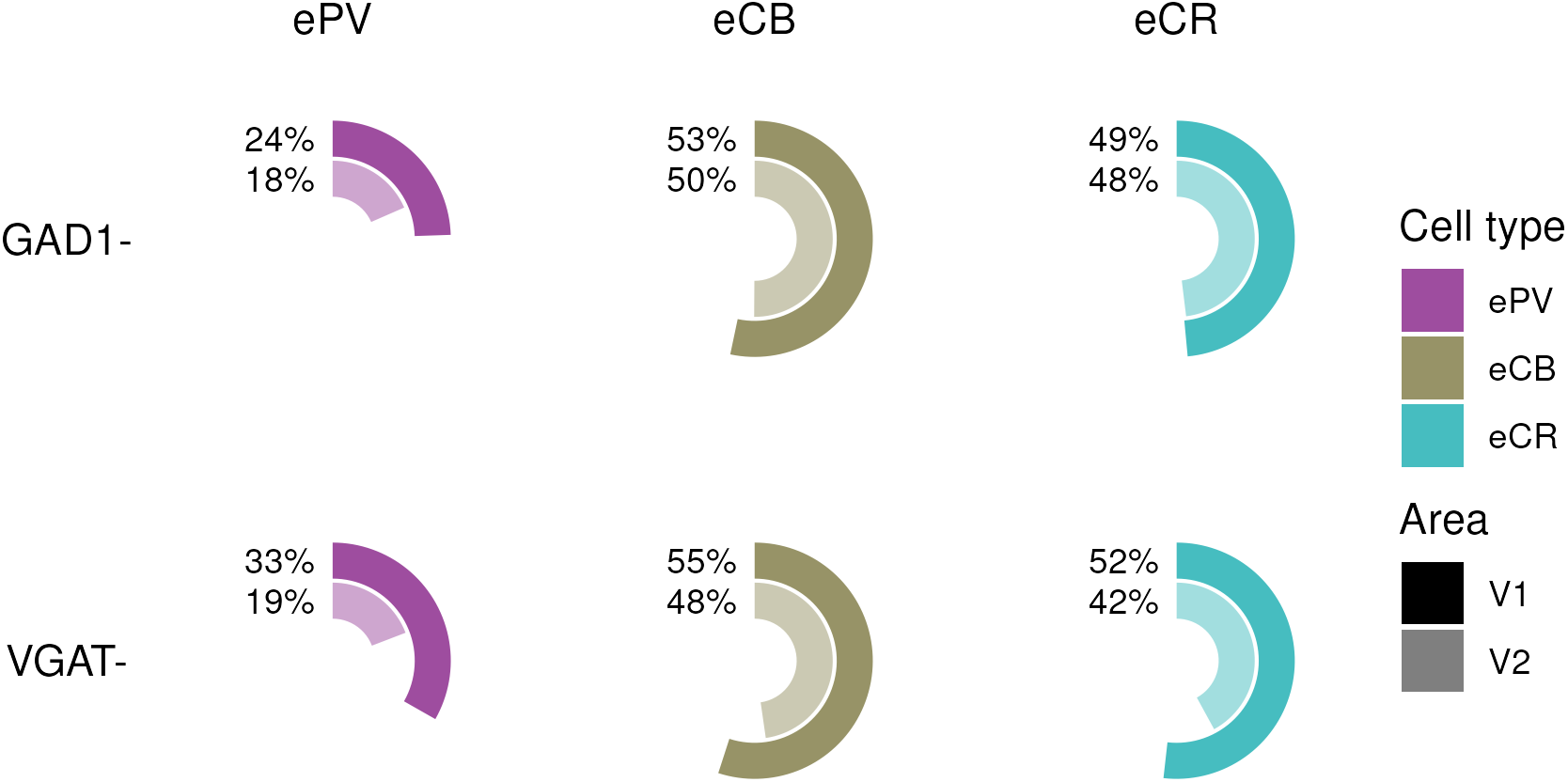
Putatively excitatory subpopulations of PV+ (ePV), CB+ (eCB), and CR+ (eCR) neurons, by 4-FISH. In each radial bar plot, the percentage of PV+ (left column), CB+ (middle), and CR+ (right) neurons that <u>do not</u> express GAD1 (GAD1-; top row) or VGAT (VGAT-; bottom) is shown for V1 (dark) and V2 (light). See **Table S2** for counts. Data from: A24.

**Table 6:** VGLUT1+ subpopulations of GAD1+ and VGAT+ neurons, by MERFISH.

| Cell type | Area | N | VGLUT1+ % (count) |
| --- | --- | --- | --- |
| GAD1+ | V1 | 44,217 | 49% (21,755) |
|  | V2 | 28,631 | 47% (13,566) |
| VGAT+ | V1 | 30,809 | 48% (14,888) |
|  | V2 | 20,077 | 46% (9,270) |

Data from: A24 (V1 and V2) and A25 (V1 only).

The proportion of non-GABAergic (GAD1-) PV neurons we observe by 4-FISH in V1 (∼25%) differs from prior reports of <10% excitatory for V1 in macaque (Kelly et al., 2019) and marmoset (Federer et al., 2024). We can reproduce the published percentages by setting a more stringent criterion on binarizing our PV count data into ‘positive’ and ‘negative’ classes; cutting off at 6 copies (5.3% GAD1-) instead of 3 (24.5% GAD1-). The higher cutoff has a correspondingly lower false discovery rate but almost certainly excludes true positives. Moving the cutoff could only reproduce the prior data if the exclusion is biased: indeed, moving the cutoff from 3 to 6 removes 87.2% of the GAD1-/PV+ population and 26.0% of the GAD1+/PV+ population.

The MERFISH detection panel we used in A25 (but not A24) includes a low-sensitivity PV probe, which we do not report elsewhere because the false negatives are too high to allow quantification. We can, however, use these data to look at what happens as the PV binarization cutoff changes. The ∼1% FPR binarization cutoff of 3 for these data yields 15.1% GAD1-/PV+ neurons (77 of 510), close to the 14.3% that 4-FISH reports at a cutoff of 4 (183 of 1,283). Of the GAD1-/PV+ neurons in the MERFISH dataset, the 72 that were also VGAT- and VGLUT1+ expressed VGLUT1 at high copy number (median 106; all ≥ 28). The median distance between a GAD1-/VGAT-/VGLUT1+/PV+ neuron and the nearest putatively inhibitory PV+ neuron (GAD1+ and/or VGAT+) was 89.9 µm, limiting likely spillover. The average number of PV copies in non-GABAergic neurons (GAD1-/VGAT-) within 20 µm of a putatively inhibitory PV+ neuron was 0.25; this dropped to 0.13 beyond 100 µm (Spearman ρ = -0.06). Correcting for spillover in the two (of 72) GAD1-/VGAT-/VGLUT1+/PV+ neurons that fall into this ‘close proximity’ group removed 1 neuron from the putatively excitatory PV population.

### Distinguishing inhibitory subpopulations of CBP neurons

Given that up to half of CBP-ir/CBP+ neurons might not be GABAergic, the question - from the standpoint of inhibitory neuron classification - becomes whether the subpopulation that <u>is</u> GABAergic can be identified anatomically. At the level of protein expression, it has been suggested that the excitatory and inhibitory subpopulations amongst CB-ir neurons can be distinguished based on staining intensity (Van Brederode et al., 1990). It is now common to apply a staining intensity counting criterion in quantitative studies of all three CBP populations in macaque, although there is no prior data supporting this approach for PV- or CR-ir neurons. For CB-ir neurons, the assertion has been that while faintly stained neurons are a heterogeneous population, intensely stained neurons are GABAergic. In our experience, however, intensely stained CB-ir neurons are not always immunoreactive for GABA (GABA-ir; **Figure 7**), and staining intensity is not obviously binary in nature, with many neurons presenting edge cases in making dark/light judgements.

**Figure 7:**
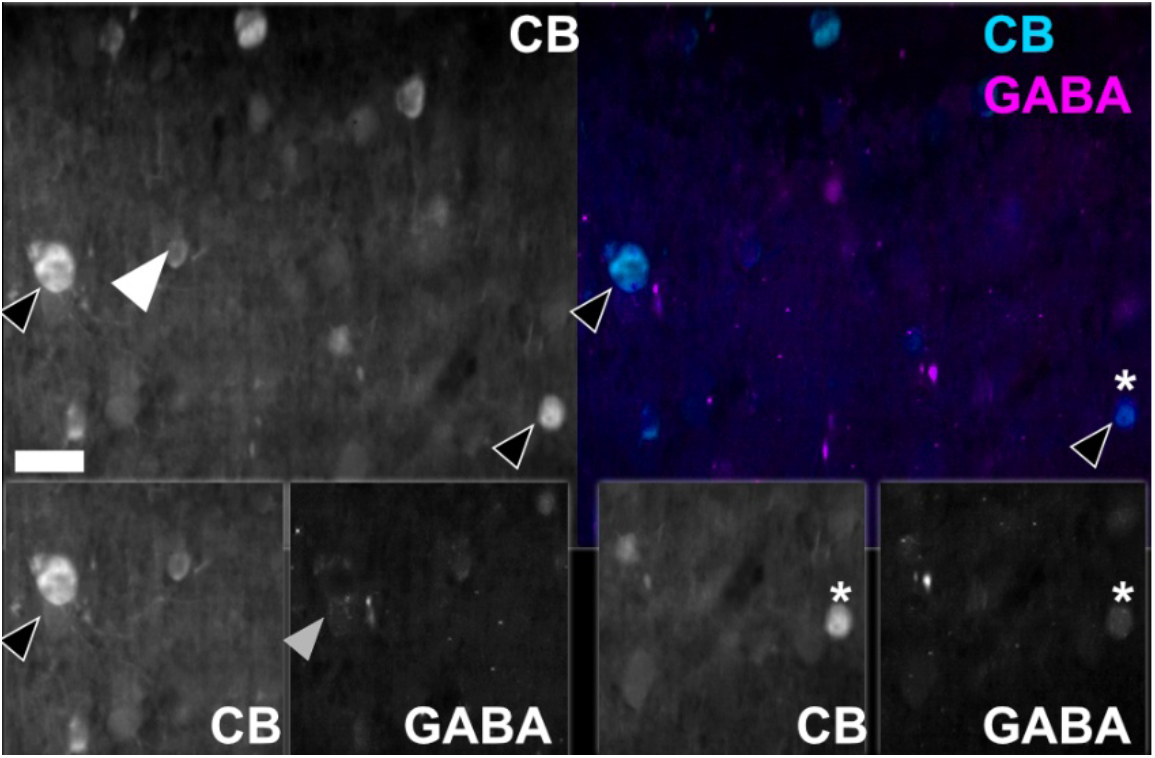
Staining intensity for CB varies from light (white arrowhead) to dark (black arrowheads). While most observers will agree on the proper categorical assignment of these particular neurons, this image shows a range of intensities that require judgment calls. This tissue was stained to visualize both CB (cyan) and GABA (magenta; upper right, dual IF; upper left, isolated CB channel). The insets below focus on the two intensely immunoreactive neurons marked by black arrowheads in the upper images. While some intensely stained CB-ir neurons are also GABA-ir (*, bottom right), others are not (bottom left, black arrowhead in isolated CB channel, gray arrowhead in isolated GABA channel). Scale bar: 20 μm. Tissue from: V1, A9.

To assess the prospect for a rigorous quantitative cutoff on fluorescence intensity for CBP immunoreactivity (regardless of its relation to what neurotransmitter is used), for each CBP we fit the normalized median intensity for every counted cell body (pooled across brain areas and animals) using Gaussian mixture modelling (GMM), with one or two Gaussian components. For both CB- and CR-ir (but not PV-ir) neurons, a two-component model was supported (Bayesian Information Criterion [BIC] difference measure, ΔBIC: CB = 114.87, CR = 417.29, PV = -87.11; **Figure 8**), although the fitted Gaussian components exhibited only moderate separation for CB (Ashman’s D = 1.55), and weak separation for CR (D = 0.95). Hartigan’s dip test, on the other hand, supported unimodality for CB and CR (p = .99 for both), and weakly supported multimodality for PV (p = .025).

**Figure 8:**
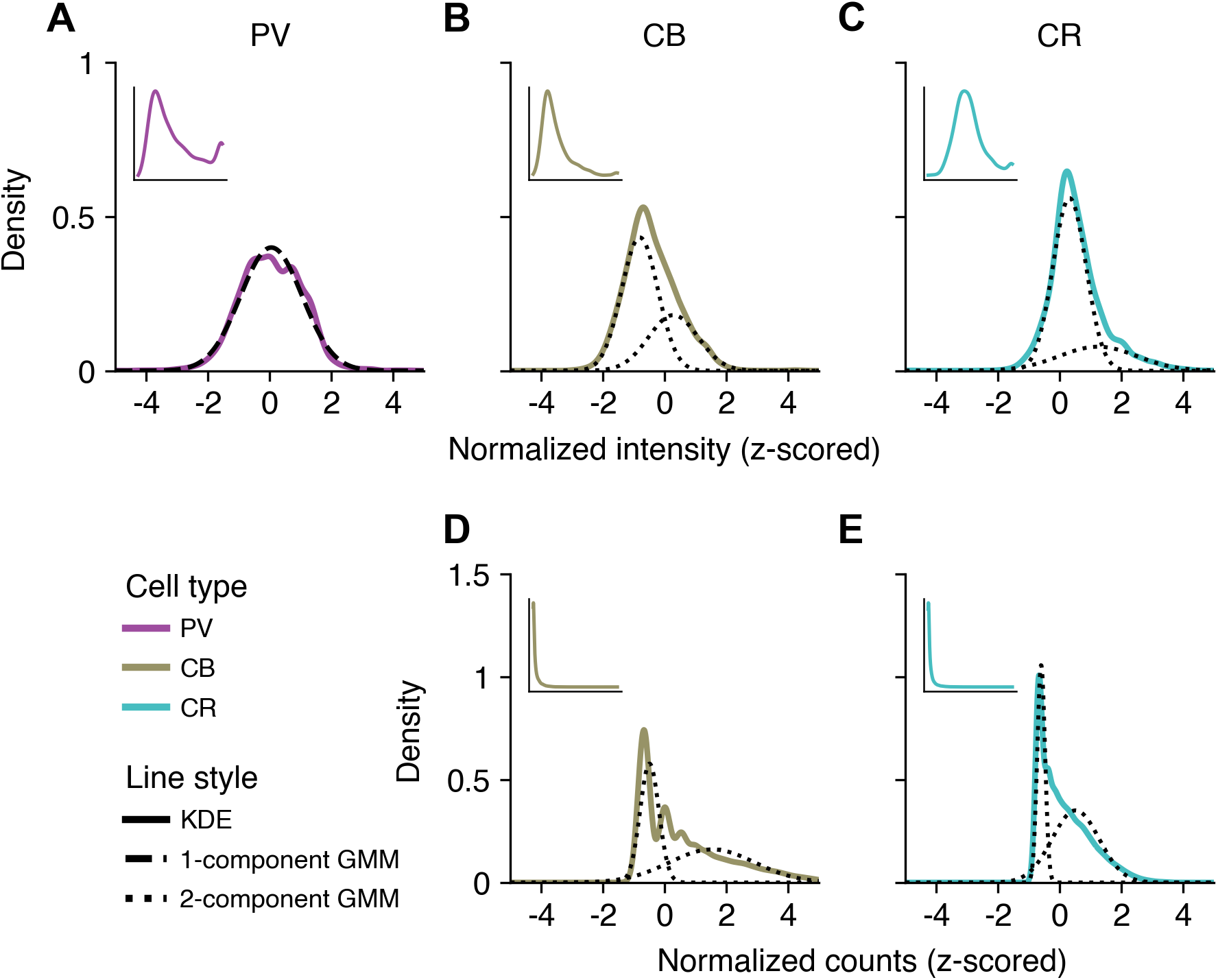
Intensity distributions of CBP-ir (A-C) and count distributions of CBP+ (D, E) neurons reveal heterogeneous but largely unimodal intensity profiles for protein expression (A-C), and multimodal count profiles for mRNA expression (D, E). Kernel density estimates (KDEs, solid lines) of normalized immunofluorescence intensity are shown for PV-ir (magenta, A), CB-ir (yellow, B), and CR-ir (cyan, C) neurons and normalized mRNA counts for CB+ (yellow, D) and CR+ (cyan, E) neurons, pooled across images. Dashed and dotted black curves show the individual mixture components for the best-fitting Gaussian mixture model (GMM: dashed, one component; dotted, two components). Insets in the upper left of each panel show KDEs of the raw (unnormalized) data. Data from: A3, A4, A8 (A-C); A24, A25 (D, E).

Proceeding despite this weak evidence for bimodality from the pooled two-component GMMs for CB-and CR-ir neurons, we constructed a model-based receiver operating characteristic (ROC) curve by sweeping the posterior threshold (area under the curve [AUC]: CB = 0.863, CR = 0.809) to establish a cross-animal intensity cutoff in normalized units. We adopted a cost-sensitive Bayes rule with a cost ratio (false positive: false negative) of 3:1, reflecting the fact that in the past, the dark/light criterion has been used to identify a population of exclusively GABAergic neurons (low false positive rate, FPR), accepting that the low intensity population is mixed (potentially high false negative rate, FNR; Van Brederode et al., 1990). Under the pooled, normalized GMMs, this corresponds to a posterior threshold *t*^\∗^ = 0.75 (z-cutoff: CB = 0.365, CR = 1.660). Applying this cutoff produces low model-based false-positive rates (FPR: CB = 0.023, CR = 0.007) at the expense of substantial false negatives (FNR: CB = 0.544, CR = 0.613) for an estimated true positive rate (TPR) of 0.456 for CB and 0.387 for CR. Performance changed minimally at cost ratios of 4:1 and 5:1, reflecting the poor peak separation of the underlying fit Gaussians. When the resulting z-cutoff (CB = 0.35, CR = 1.640) is applied to the normalized median intensity data, the fraction of cells labeled “high intensity” (i.e. putatively high-confidence GABAergic neurons) is 17.4% for CB and 10.5% for CR. Comparing these values to the above estimation of the putatively excitatory population (**Figure 6**) suggests that this approach - at best - captures about one half of true positives for GABAergic CB-ir neurons, and about one fifth for GABAergic CR-ir neurons, assuming intensely stained neurons are, in fact, GABAergic.

While intense protein immunoreactivity does not necessarily arise from an underlying high mRNA copy count, we nonetheless made use of data on-hand to run an equivalent analysis on mRNA. One- and two-component GMMs were fit to copy count data for 44,951 cells with positive probe hybridization for CB and/or CR (2,698 cells that were CB+/CR+ were included in both populations) from the MERFISH experiments in A24 and A25 (the PV probe used in these experiments is not analyzed, see Methods). In both cases, a two-component model was preferred (ΔBIC: CB = 34,711, CR = 34,378) and peak separation was moderate (Ashman’s D: CB = 1.42, CR = 1.26).

Unlike in the above IF study, GAD1 and VGAT were probed in the MERFISH panel, so we have ground truth. ROC AUCs were similar whether we defined ‘GABAergic’ neurons as those expressing mRNA for GAD1 only (AUC: CB = 0.635, CR = 0.697), VGAT only (AUC: CB = 0.668, CR = 0.691), or both GAD1 and VGAT (AUC: CB = 0.672, CR = 0.700). Sensitivity was best for GAD1+/VGAT+ neurons (CB: TPR = 0.403, FPR = 0.144, FNR = 0.597; CR: TPR = 0.402, FPR = 0.102, FNR = 0.598) and worst for GAD1 only (CB: TPR = 0.346, FPR = 0.144, FNR = 0.654; CR: TPR = 0.330, FPR = 0.047, FNR = 0.670). Classification performance was identical for FP:FN cost ratios of 3:1, 4:1, and 5:1. Thus, on a ground truth dataset, classification performance for mRNA count data (TPR ∼0.33-0.40; FPR ∼0.05-0.15) was similar to model-based performance on protein intensity data, with a notably higher false positive rate for mRNA.

### Criterion 3: A substantial fraction of GABAergic neurons does not express parvalbumin, calbindin, or calretinin

We found that 52% of GAD1+ cells in V1 were PV+ (1,207 of 2,335; 4-FISH, A24 only, slides 18 and 19, see **Table 4** for 4-FISH experiment design), 13% were CB+ (6,041 of 46,472; 4-FISH and MERFISH, A24 and A25), and 15% were CR+ (6,846 of 46,472; 4-FISH and MERFISH, A24 and A25). This is less than the total GABAergic population; accordingly, 25% of GAD1+ cells (221 of 900; 4-FISH, A24, slide 19) expressed no CBP mRNA (**Figure 3, top**). Due to co-expression, these values sum to 105%. In V2, the percentages were 38% PV+ (380 of 1,009), 14% CB+ (4,316 of 29,891), 26% CR+ (7,683 of 29,891), and 33% CBP- (149 of 453). The result was similar for VGAT+ cells. As it does in mouse cortex (Rudy et al., 2011), the alternate classification scheme that uses PV, SST, and VIP left a similarly-sized population of GAD1+ cells unaccounted for (4-FISH only, V1: 321 of 1,435 GAD1+ neurons, 22%, **Figure 3, bottom**; V2: 166/556, 30%). Again, the result was similar for VGAT+ cells (V1: 145 of 838 VGAT+ neurons, 17%, **Figure S4, bottom**; V2: 87/367, 24%). Co-expression of PV/VIP/SST is as rare as co-expression of PV/CB/CR (<5% for any combination; 4-FISH **Table S5**).

### Criterion 4: Neurons immunoreactive for parvalbumin, calbindin, and calretinin can be roughly mapped to the parvalbumin/somatostatin/vasoactive intestinal peptide scheme used in other species

We probed mRNA expression of VIP and SST in CB+ and CR+ neurons by 4-FISH and MERFISH and found that CB+ neurons in V1 and V2 rarely express mRNA for VIP, and CR+ neurons rarely express mRNA for SST (4-FISH only: **Tables 7-8**; raw data, **Figure S6**; laminar co-expression with PV, **Table S6**; 4-FISH and MERFISH combined: plot of co-expression by layer, **Figure S7**; laminar count data: **Tables S7, S8**). Two neurons in V2 that expressed mRNA for <u>both</u> CB and CR were SST+. The CBP+ and peptide mRNA-expressing populations are not identical, however: many neurons express mRNA for a CBP, or a peptide, but not both.

**Table 7:** Expression of SST and VIP by CB+ and CR+ neurons, by 4-FISH.

| Area | CBP cell type | N | SST+ % (count) | VIP+ % (count) |
| --- | --- | --- | --- | --- |
| V1 | CB+ | 252 | 62% (155) | 2% (4) |
|  | CR+ | 328 | 5% (18) | 62% (202) |
| V2 | CB+ | 169 | 54% (92) | 1% (2) |
|  | CR+ | 293 | 8% (24) | 51% (149) |

Data from: A24.

**Table 8:**
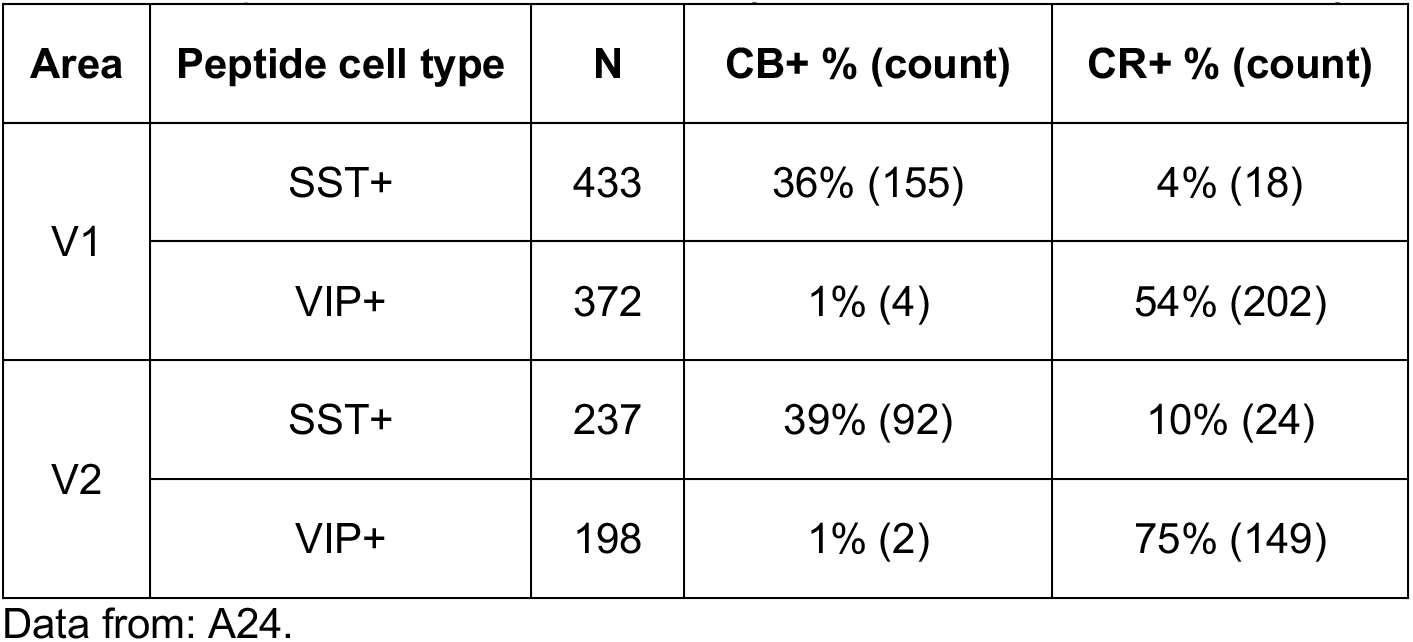
Expression of CB and CR by SST+ and VIP+ neurons, by 4-FISH.

| Area | Peptide cell type | N | CB+ % (count) | CR+ % (count) |
| --- | --- | --- | --- | --- |
| V1 | SST+ | 433 | 36% (155) | 4% (18) |
|  | VIP+ | 372 | 1% (4) | 54% (202) |
| V2 | SST+ | 237 | 39% (92) | 10% (24) |
|  | VIP+ | 198 | 1% (2) | 75% (149) |

Data from: A24.

Interestingly, in MERFISH experiments that allowed us to probe for GAD1, VGAT, CB, CR, SST, and VIP (**Figure 9**) in the same tissue section, the vast majority of neurons that express mRNA for <u>both</u> a peptide and a CBP were inhibitory (**Table 9**). Detection of VIP by MERFISH was poor (low copy count), raising concerns about false negatives in this experiment. In a follow-up experiment by 4-FISH, we observed similar proportions of putatively inhibitory neurons amongst the VIP+ population as in the MERFISH experiment, whether the marker was VGAT or GAD1 (4-FISH only: **Table S9**, counts; **Figure 3, bottom**, GAD1 raw data; **Figure S4**, bottom, VGAT raw data; 4-FISH and MERFISH combined: **Figure S7**, laminar co-expression). It was still the case that VIP+ neurons were more likely to be putatively inhibitory if they also expressed CR, for both VGAT and GAD1 (**Table S9**). SST and VIP alone do a better job of identifying inhibitory neurons (being 62-88% inhibitory depending on GABA marker) than do CB and CR (42-70% inhibitory; **Table 9**).

**Figure 9:**
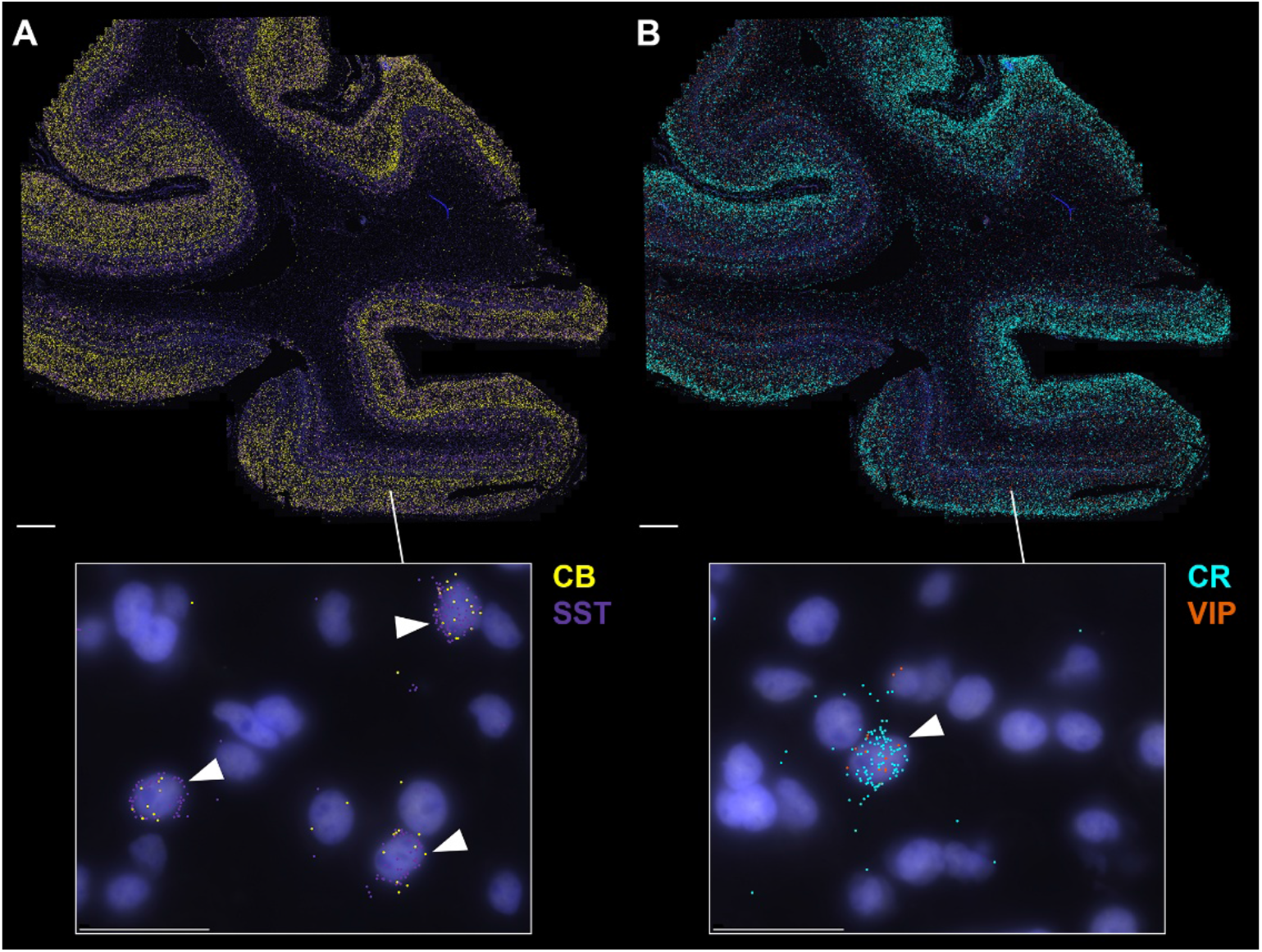
Spatial transcriptomic data (MERFISH) for CBP and neuropeptide mRNA transcripts in early visual cortex. Both panels show the same tissue section with DAPI (blue) and PolyT (white) staining and colored dots representing mRNA transcript locations for CB (yellow, A), SST (purple, A), CR (cyan, B), and VIP (orange, B) in cortical areas V1 and V2. Insets show double-labeled cells (white arrows) from layers 2 and 3 of V1 with co-localization of CB/SST (A) and CR/VIP (B). Data from: A24. Scale bar: 1 mm, panels; 25 μm, insets.

**Table 9:** Putatively inhibitory subpopulations of SST+, VIP+, CB+, CR+, SST+/CB+, and VIP+/CR+ neurons, by MERFISH.

| Cell type | Area | N | GAD1+ % (count) | VGAT+ % (count) |
| --- | --- | --- | --- | --- |
| SST+ | V1 | 11,125 | 82% (9,082) | 63% (6,976) |
|  | V2 | 8,778 | 82% (7,239) | 62% (5,486) |
| VIP+ | V1 | 3,180 | 85% (2,700) | 67% (2,126) |
|  | V2 | 2,629 | 88% (2,306) | 66% (1,727) |
| CB+ | V1 | 10,875 | 51% (5,561) | 42% (4,571) |
|  | V2 | 5,921 | 69% (4,077) | 59% (3,494) |
| CR+ | V1 | 9,317 | 68% (6,379) | 50% (4,627) |
|  | V2 | 10,483 | 70% (7,322) | 47% (4,950) |
| SST+/CB+ | V1 | 3,860 | 94% (3,647) | 80% (3,104) |
|  | V2 | 3,036 | 97% (2,931) | 83% (2,521) |
| VIP+/CR+ | V1 | 2,602 | 93% (2,411) | 74% (1,919) |
|  | V2 | 2,334 | 93% (2,171) | 70% (1,628) |

Data from: A24 (V1 and V2) and A25 (V1 only).

## Discussion

We set out to understand the extent to which the long-standing practice of using immunoreactivity for calcium-binding proteins (CBPs) to classify GABAergic neurons in macaque cortex is justified, and conclude it is not a strong approach without a secondary GABAergic cell marker. Suitable alternative schemes include PV, CB/SST, and CR/VIP, or simply PV/SST/VIP. While it remains possible that the CBP-based scheme can be used validly in cortical areas we did not study, it should not be deployed in these areas without prior validation.

Our counts were non-stereological; appropriate for studying proportions and co-expression, but not for estimating density or total number. This was the right approach here; stereological data become relevant once a ‘parts list’ is established, and a core goal of this work was to contribute to determining what that parts list should be for macaque cortex. A key concern with non-stereological approaches is size bias: Abercrombie correction did not change overall proportions, indicating that cell type-specific differences in soma sizes/aspect ratios did not bias our results.

### How should inhibitory neurons be classified in macaque cortex?

To guide our decisions, we set criteria on validity and utility.

#### Criterion 1 is met

We find that > 95% of CBP-ir (protein-expressing) neurons express only one CBP, allowing them to be assigned to a single class; whether class membership would change over time in adult cortex is not known. All prior studies agree that CBP co-expression is minimal in macaque cortex (Van Brederode et al., 1990; Kooijmans et al., 2020; Medalla et al., 2023), in contrast to other species (Kawaguchi and Kubota, 1997; Hof et al., 1999).

#### Criterion 2 is not met

We find that substantial fractions of CBP+ (mRNA-expressing) neurons are not GABAergic, making the identification of GABAergic subpopulations critical. Our finding that only ∼75% of PV+ neurons in V1 are GABAergic is similar to our own prior study (Constantinople et al., 2009) and differs from Kelly et al. (2019). We do, however, replicate their inverse calculation: 52% of GABAergic neurons express PV (not the 74% reported by Van Brederode et al., 1990). A failure of GABAergic neuron detection is unlikely: in our study, the proportion of segmented DAPI nuclei that were GAD1 positive was very similar across two mRNA detection methods: 4-FISH 12% and MERFISH 13%. These GAD1+ neurons comprised 20% of the neuronal population (72,848 of 364,408) in the MERFISH study, which comports well will a prior study that detected GABA directly (Beaulieu et al., 1992).

We attribute the difference between our estimation of the excitatory PV population and Kelly et al.’s to the threshold at which a neuron is scored PV+: we found that non-GABAergic PV+ neurons express PV at low copy number and that restricting our counts to the highest-expressing cells approximately reproduces the Kelly et al. result. The neurons excluded by this more stringent cutoff were strongly biased towards the GAD1-population. The neurons above the cutoff strongly express VGLUT1, lack GAD1 and VGAT, and cannot be accounted for by spillover from neighboring PV+/GAD1+ neurons. A similar threshold difference, rather than a species difference we cannot formally exclude, may likewise account for the high GABAergic fraction reported in marmoset V1 by Federer et al. (2024). Our findings agree well with the prior report that CB-ir neurons comprise 12% (we find 13%) of the GABAergic population (Van Brederode et al., 1990) and with the 14% of GABAergic neurons reported to be CR-ir (we find 15%; Meskenaite, 1997).

Intensity criteria are not supported for CB- or CR-ir neurons; identifying inhibitory subpopulations within these classes requires a GABAergic cell marker. mRNA detection would work, but does not yield morphological information and, in our experience, false negatives arise when *in situ* hybridization for mRNA is combined with immunohistochemistry for peptides/proteins. Immunohistochemical visualization of GABAergic neurons also presents challenges: antibodies against GAD yield false negatives without colchicine, and antibodies against GABA require glutaraldehyde fixation, which causes false negatives for CBPs.

Our mRNA data support co-labeling for CB/SST and CR/VIP, which yields populations that are >90% inhibitory. This arises because VIP+ and SST+ populations are largely inhibitory, so it is also probably appropriate to adopt PV/SST/VIP classification. The PV+, SST+, and VIP+ populations were non-overlapping (criterion 1), and this scheme left a similar fraction of GABAergic neurons unaccounted for as did PV/CB/CR (criterion 3). An important caveat: while putatively inhibitory CB+ neurons in macaque cortex express VIP, they do not express any type-3 serotonin receptors (5-HT3R; unpublished data) and so are <u>not</u> functionally identical to VIP+ neurons in mice. This implies that classification using PV/SST/5-HT3R expression, which in mice accounts for ∼100% of GABAergic neurons (Rudy et al., 2011), will not port to macaque. Furthermore, the ‘canonical’ cortical motif often invoked to describe connectivity between excitatory neurons and PV, SST, and VIP inhibitory neurons in the superficial layers of mouse cortex (Pfeffer et al., 2013) will probably play out differently across layers in macaque cortex, because SST and VIP neurons (subsets of CB and CR neurons) are rarely found in layers 4-6. Study of the synaptic circuitry of excitatory-inhibitory interactions in these deeper layers seems warranted.

### Inter-area differences with a conserved laminar motif

With particularly high PV and low CR counts, V1 is the ‘oddball’ amongst the areas we studied. Macaque V1 is anatomically distinct - with respect to V1 in other species, and other areas in macaque - in numerous ways, including packing density, proportion of inhibitory neurons (Rockel et al., 1980; Collins et al., 2010; Turner et al., 2016; Kelly et al., 2019), and laminar organization (Casagrande and Kaas, 1994; Brodmann, 2005). The population composition we report for V1 differs substantially from Medalla et al. (Medalla et al., 2023) and less so from Kooijmans et al. (Kooijmans et al., 2020), probably because we all used different counting criteria. Medalla et al. counted “strongly immunolabeled neurons” with non-pyramidal morphology for all classes; Kooijmans et al. applied an intensity criterion to CB-ir counts only; we applied no intensity criterion for any type. It is unsurprising that, having counted different things, our results differ.

V2 and V3 have strikingly similar compositions, perhaps supporting an ‘early extrastriate’ phenotype, when compared with V3A. Amidst ongoing disagreement about the organization of ‘third-tier’ visual cortex in primates (Angelucci and Rosa, 2015; Arcaro and Kastner, 2015; Gattass et al., 2015; Kaas et al., 2015), our data support anatomically distinct tissue in the dorsal cortex anterior/medial to V3. Beyond this, V3A, V4, and MT are compositionally distinct, with large PV-ir populations. By prefrontal cortex, CR-ir neurons predominate (Condé et al., 1994; Medalla et al., 2023); what happens in between is unknown.

### Functional interpretability of cell type classification schemes

It is now common to classify neurons using transcriptomic markers that can be difficult to interpret functionally. VIP and SST, however, are bioactive peptides, expression of which can support mechanistic prediction. Stored in dense core vesicles, SST and VIP are probably co-released under high firing rates, yielding a second signaling regime: GABA alone versus GABA plus peptide. VIP binds VIP/PACAP family receptors coupled to Gs and Gq, so the secondary regime is likely ‘excitatory’. SST receptors (SST1-5) are all Gi/o coupled, implying an ‘inhibitory’ secondary regime. Peptide receptors are differentially expressed across cortical neurons (Smith et al., 2019), suggesting partner-specific effects. Thus, layering the functional consequences of peptide expression over traditional circuit motifs should yield nontrivial predictions.

Similarly, CBP expression likely has implications for plasticity. PV has a low molecular weight and lacks protein-protein interaction sites that could limit diffusion. If the readily releasable vesicle pool can keep up, the resulting high-capacity, mobile buffering can support fast neuronal firing (Kawaguchi et al., 1987; Müller et al., 2007). If the pool <u>cannot</u> keep up, synapses might depress, but are unlikely to facilitate. CB and CR, in contrast, both interact with other proteins (Schwaller et al., 2002; Schwaller, 2009), allowing anchoring, trafficking, modulation, and sensor-like behavior (not just buffering). CR’s binding sites show strong cooperativity, producing a nonlinear affinity that increases with calcium concentration and so likely also limits facilitation. A fast buffer, CB can clip presynaptic calcium transients, helping delay or prevent depression, but at some synapses, CB saturation can lead to ‘pseudofaciliation’ (Neher, 1998; Rozov et al., 2001; Schmidt, 2012; Jackman and Regehr, 2017). Postsynaptically, these buffering characteristics likely yield different history dependence for the calcium transients that drive many forms of longer-term plasticity.

### Concluding remark

Beyond any “parts lists”, knowing which peptides and CBPs a neuron expresses can support functional prediction. The proper cell typing scheme then depends on the question(s) under study and - perhaps more importantly, and neglected - can be used not only to infer synaptic partners but also to predict how a neuron will interact with those partners. In the end, this is what neurons are using these ‘markers’ for: not as nametags, but as functional elements.

## Financial Interests

The authors declare no competing interests.

## Acknowledgements and Author Contributions

This work was supported by NIH grants R00MH093567 and R01EY029663 **to AAD** and NSF Graduate Research Fellowship DGE 2139754 **to AMB**. The authors gratefully acknowledge the support of M. Yankoswki and H. Alasady for assistance with histological processing and antibody controls. AAD designed and supervised the experiments, processed tissue for the immunohistochemical and immunofluorescence analyses, and collected and analyzed immunofluorescence raw data. AMB undertook the *in situ* hybridization studies and designed and executed the machine learning approach to fluorescence intensity. JK and CP helped analyze immunofluorescence data. AAD, AMB, and JK co-wrote the manuscript. Opinions, findings, and conclusions or recommendations expressed in this material are those of the author(s) and do not necessarily reflect the views of the National Science Foundation or the National Institutes of Health.

## Notes

### Competing Interest Statement

The authors have declared no competing interest.

### Summary of Updates

Update author affilitation, add data. No changes to findings.

