## Supplementary Information for "Calcium-binding protein expression alone is insufficient to identify and classify GABAergic neurons in macaque cortex"

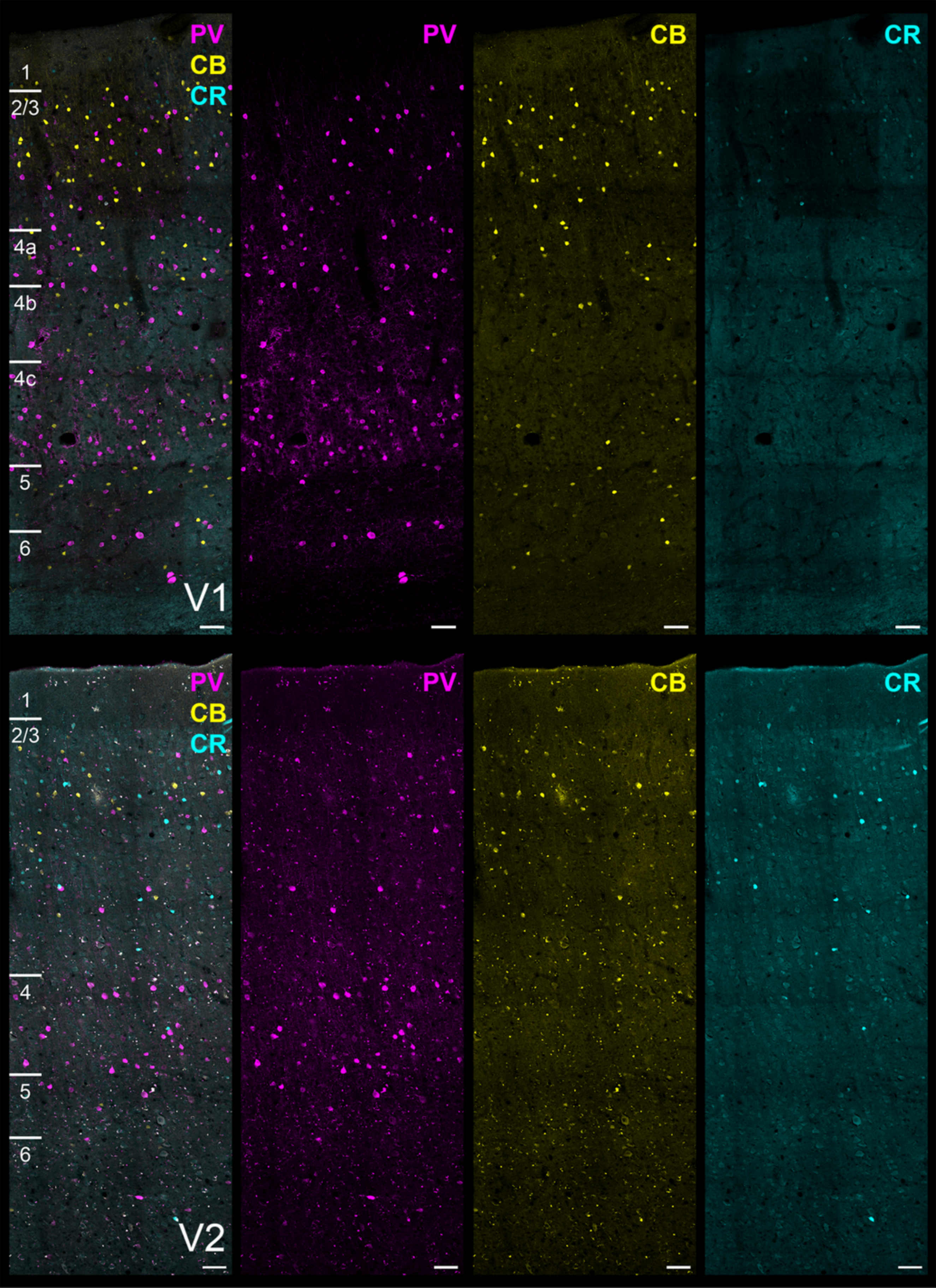
PV
CB
CR
1
2/3
4a
4b
4c
5
6
V1
PV
CB
CR
PV
CB
CR
1
2/3
4
5
6
V2
PV
CB
CR

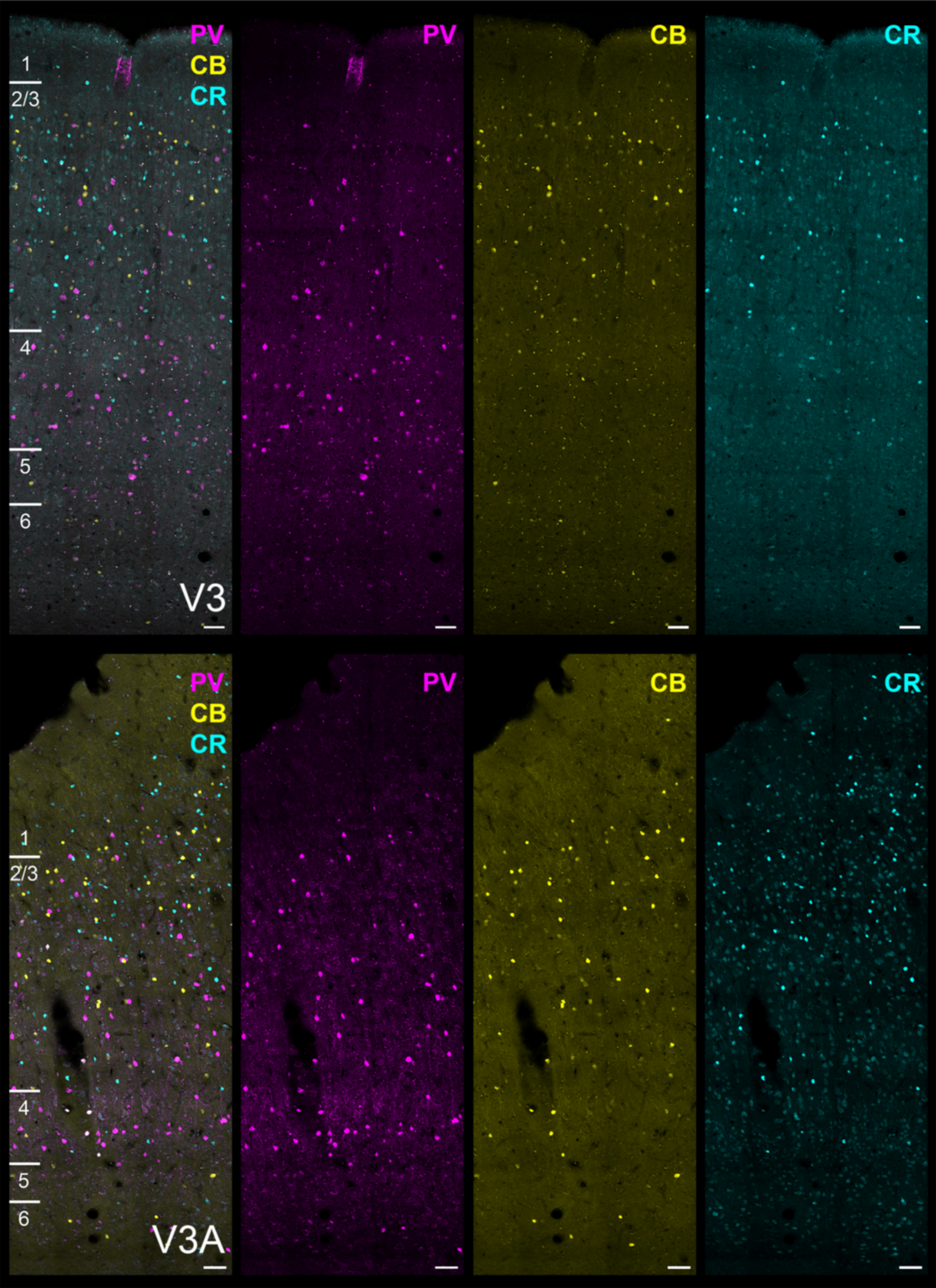
PV
CB
CR
1
2/3
4
5
6
V3
PV
CB
CR
PV
CB
CR
1
2/3
4
5
6
V3A
PV
CB
CR

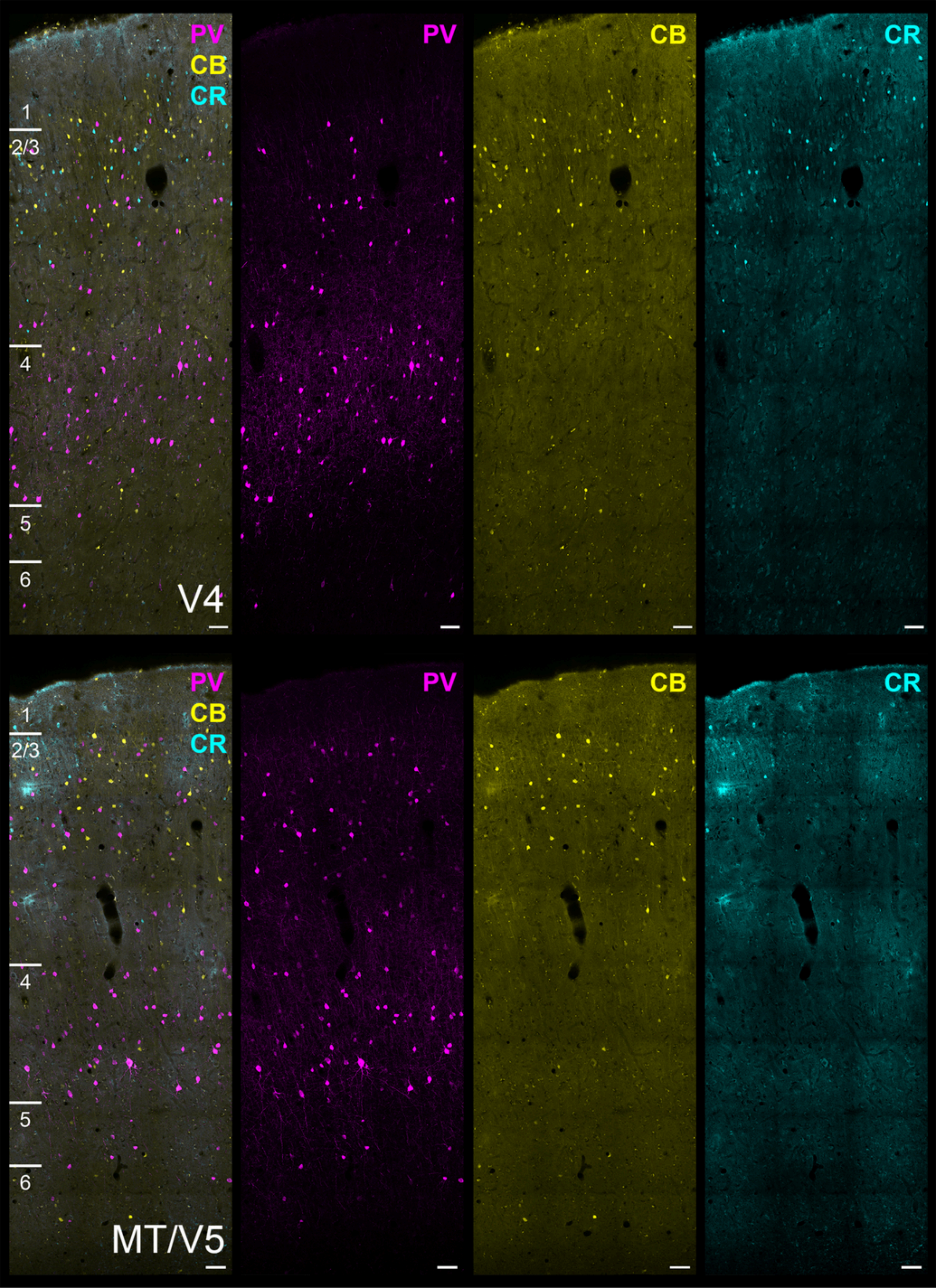

PV
CB
CR
1
2/3
4
5
6
V4
PV
CB
CR
PV
CB
CR
1
2/3
4
5
6
MT/V5
PV
CB
CR

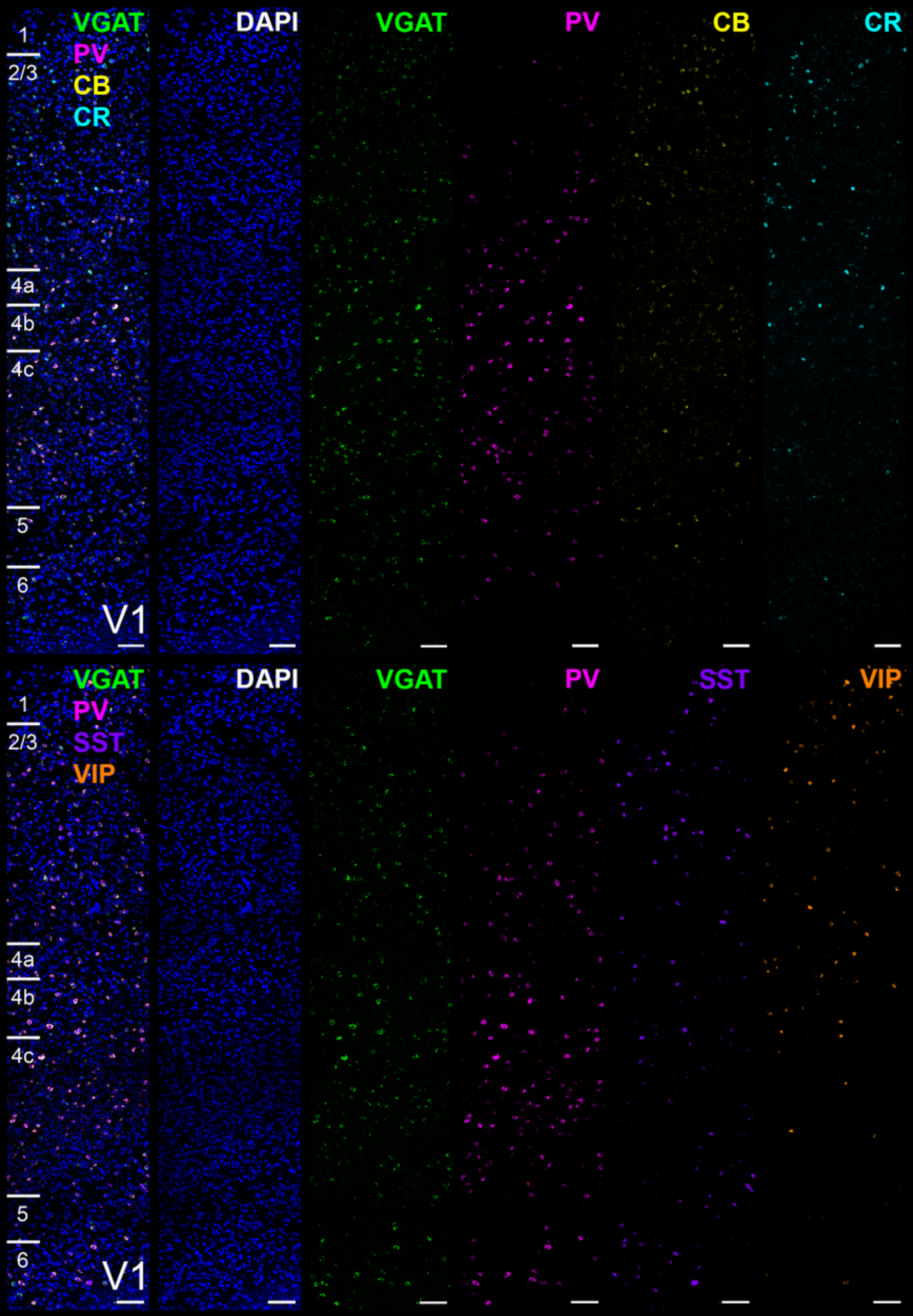

VGAT
PV
CB
CR
1
2/3
4a
4b
4c
5
6
V1
DAPI
VGAT
PV
CB
CR
VGAT
PV
SST
VIP
1
2/3
4a
4b
4c
5
6
V1
DAPI
VGAT
PV
SST
VIP

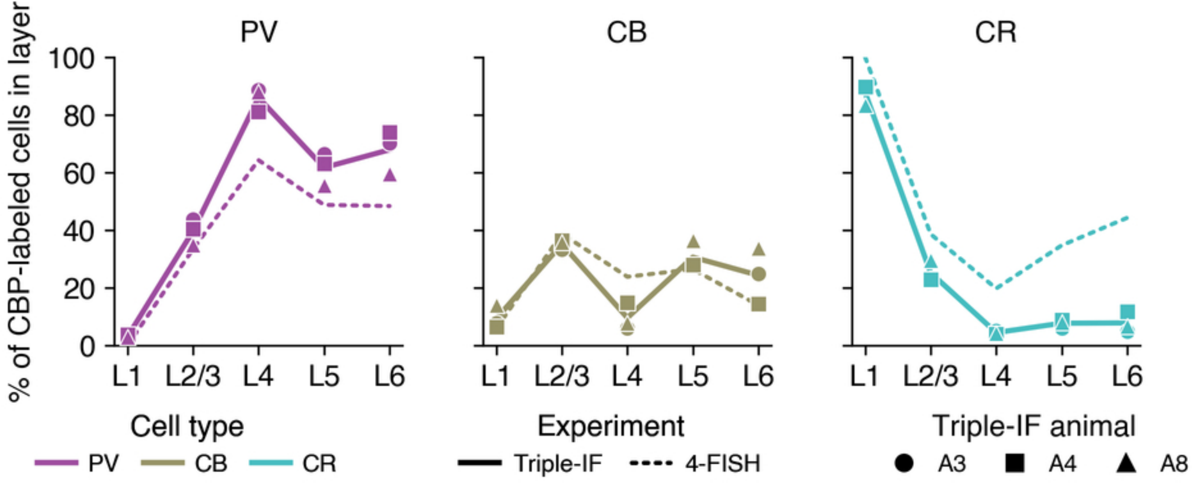
PV
CB
CR
% of CBP-labeled cells in layer
100
80
60
40
20
0
L1
L2/3
L4
L5
L6
Cell type
PV
CB
CR
Experiment
Triple-IF
4-FISH
Triple-IF animal
A3
A4
A8

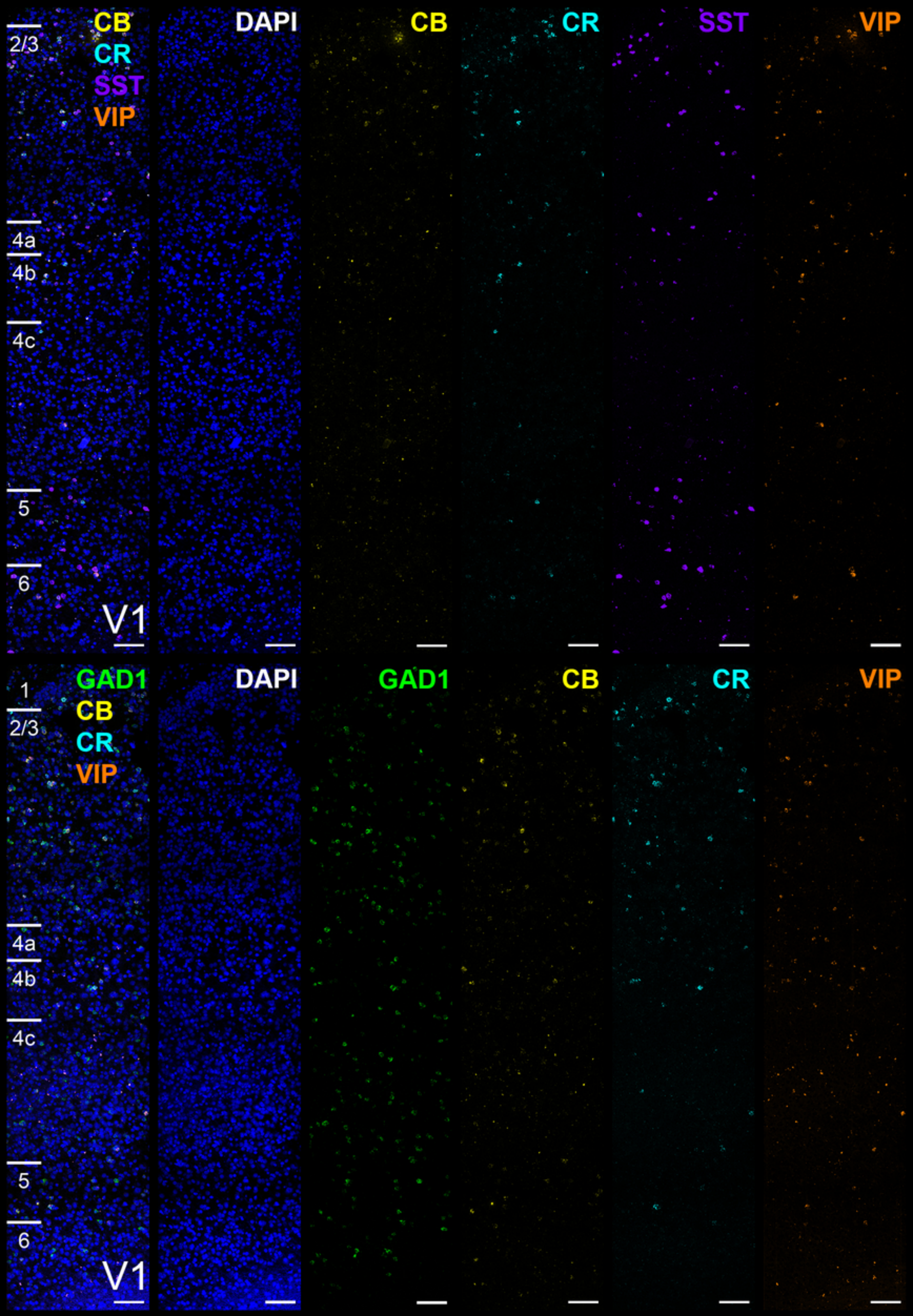

CB
CR
SST
VIP
DAPI
CB
CR
SST
VIP
2/3
4a
4b
4c
5
6
V1
GAD1
CB
CR
VIP
DAPI
GAD1
CB
CR
VIP
1
2/3
4a
4b
4c
5
6
V1

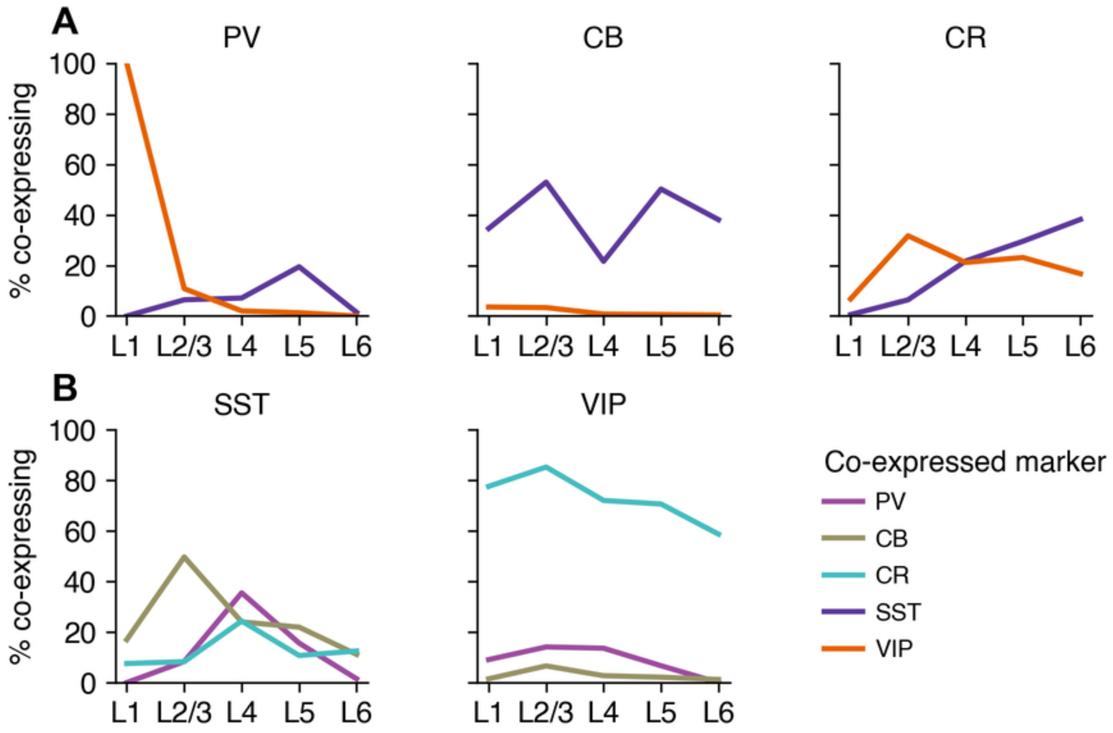
A
PV
CB
CR
100
80
60
40
20
0
% co-expressing
L1
L2/3
L4
L5
L6
B
SST
VIP
Co-expressed marker
PV
CB
CR
SST
VIP

### Supplemental Figure Legends

**Figure S1:** Triple Immunofluorescence labeling for CBPs in early visual cortex. Immunofluorescence is shown for PV (magenta), CB (yellow) and CR (cyan) in cortical areas V1 (top row) and V2 (bottom row). Cortical layer borders are indicated to the left of each panel; note that layer boundaries in the tissue are not parallel with image borders/layer markers in all panels. Data from: A4 (V1), A8 (V2). Scale bar: 50 µm, all panels.

**Figure S2:** Triple Immunofluorescence labeling for CBPs in early- to mid-level visual cortex. Immunofluorescence is shown for PV (magenta), CB (yellow) and CR (cyan) in cortical areas V3 (top row) and V3A (bottom row). Cortical layer borders are indicated to the left of each panel; note that layer boundaries in the tissue are not parallel with image borders/layer markers in all panels. Data from: A4 (V3A), A8 (V3). Scale bar: 50 µm, all panels.

**Figure S3:** Triple Immunofluorescence labeling for CBPs in mid-level visual cortex. Immunofluorescence is shown for PV (magenta), CB (yellow) and CR (cyan) in cortical areas V4 (top row) and MT/V5 (bottom row). Cortical layer borders are indicated to the left of each panel; note that layer boundaries in the tissue are not parallel with image borders/layer markers in all panels. Data from: A3. Scale bar: 50 µm, all panels.

**Figure S4:** 4-plex RNAscope fluorescent labeling (4-FISH) for CBP (top row) and neuropeptide (bottom row) mRNA transcripts in V1. DAPI-stained nuclei (blue) with fluorescent puncta (mRNA transcripts) are shown for VGAT (green), PV (magenta), CB (yellow), CR (cyan), SST (purple), and VIP (orange). Cortical layer borders are indicated to the left of each panel; note that layer boundaries in the tissue are not parallel with image borders/layer markers in all panels. Experimental design in **Table 4**. Data from: A24. Scale bar: 100 µm, all panels.

**Figure S5:** Comparison of CBP-ir (protein) population composition and CBP+ (mRNA; 4-FISH) population compositions, by layer in V1 and V2. Comparison of PV (magenta; left), CB (yellow, middle), and CR (cyan, right) expressing neurons as a percentage of all neurons in each layer. Composition is similar whether measured by  protein (Triple-IF, solid lines and individual animal markers) versus mRNA (4-FISH, dashed line, one animal only). Data for layers 4a and 4b of V1 excluded. Data from: A3, A4, A8 (Triple-IF) and A24 (4-FISH).

**Figure S6:** 4-plex RNAscope fluorescent labeling (4-FISH) for CBP and neuropeptide mRNA transcripts in V1. DAPI-stained nuclei (blue) with fluorescent puncta (mRNA transcripts) are shown for GAD1 (green), PV (magenta), CB (yellow), CR (cyan), SST (purple), and VIP (orange) in cortical area V1. Cortical layer borders are indicated to the left of each panel; note that layer boundaries in the tissue are not parallel with image borders/layer markers in all panels. Experimental design in **Table 4**. Data from: A24. Scale bar: 100 µm, all panels.

**Figure S7:** Laminar co-expression of CBPs with SST and VIP, by layer in V1 and V2, by 4-FISH and MERFISH. Top row: Percentage of PV+ (left), CB+ (middle), and CR+ (right) neurons that also express SST (purple) and or VIP (orange), by layer, pooled across V1 and V2. Bottom row: Percentage of SST (left) and VIP (middle) neurons that also express PV (magenta), CB (yellow), or CR (cyan), by layer, pooled across V1 and V2. Data for layers 4a and 4b of V1 excluded. Data from: A24 and A25 (4-FISH and MERFISH).

**Supplemental Tables**

**Table S1:** Laminar data for PV, CB, and CR in V1 and V2, by Triple-IF versus 4-FISH.

| CBP | Layer | % of total CBP-labeled cells in layer, Triple-IF (count) | % of total CBP-labeled cells in layer, 4-FISH (count) |
|---|---|---|---|
| PV | 1 | 3% (5 of 154) | 0% (0 of 17) |
| | 2/3 | 40% (3,061 of 7,692) | 33% (488 of 1,461) |
| | 4a (V1 only) | 71% (10 of 14) | 75% (61 of 81) |
| | 4b (V1 only) | 82% (45 of 55) | 50% (102 of 205) |
| | 4/4c | 86% (1,678 of 1,952) | 64% (853 of 1,327) |
| | 5 | 63% (571 of 911) | 49% (59 of 121) |
| | 6 | 70% (717 of 1,020 | 48% (59 of 122) |
| CB | 1 | 8% (13 of 154) | 6% (1 of 17) |
| | 2/3 | 35% (2,719 of 7,692) | 39% (566 of 1,461) |
| | 4a (V1 only) | 29% (4 of 14) | 28% (23 of 81) |
| | 4b (V1 only) | 15% (8 of 55) | 41% (84 of 205) |
| | 4/4c | 10% (186 of 1,952) | 24% (316 of 1,327) |
| | 5 | 30% (269 of 911) | 15% (32 of 121) |
| | 6 | 21% (213 of 1,020) | 14% (17 of 122) |
| CR | 1 | 88% (136 of 154) | 100% (17 of 17) |
| | 2/3 | 25% (1,912 of 7,692) | 39% (564 of 1,461) |

| | | | |
|---|---|---|---|
| | 4a (V1 only) | 0% (0 of 14) | 19% (15 of 81) |
| | 4b (V1 only) | 15% (8 of 55) | 21% (43 of 205) |
| | 4/4c | 5% (88 of 1,952) | 20% (262 of 1,327) |
| | 5 | 8% (71 of 911) | 35% (42 of 121) |
| | 6 | 9% (90 of 1,020 | 44% (54 of 122) |

Triple-IF data from: A3, A4, A8; 4-FISH data from: A24. Percentages per layer sum to >100 (particularly in 4-FISH) due to co-expression.

**Table S2:** Putatively excitatory subpopulations for PV+ (ePV), CB+ (eCB), and CR+ (eCR) neurons, by 4-FISH.

| GABA marker | Area | ePV % (count) | eCB % (count) | eCR % (count) |
|---|---|---|---|---|
| GAD1- | V1 | 24% (391 of 1,598) | 53% (548 of 1,028) | 49% (440 of 907) |
| | V2 | 18% (86 of 466) | 50% (240 of 479) | 48% (334 of 695) |
| VGAT- | V1 | 33% (423 of 1,273) | 55% (403 of 733) | 52% (247 of 477) |
| | V2 | 19% (65 of 339) | 48% (303 of 635) | 42% (290 of 690) |

Data from: A24.

**Table S3**: Laminar distribution of putatively excitatory PV neurons (ePV) in V1 and V2, by 4-FISH.

| CBP | Layer | % ePV by GAD1 (count) | % ePV by VGAT (count) |
|---|---|---|---|
| PV | 1 | 0% (0 of 1) | n/a |
| | 2/3 | 19% (110 of 591) | 23% (137 of 599) |
| | 4/4c | 28% (283 of 1,004) | 38% (264 of 693) |
| | 5 | 9% (7 of 80) | 30% (17 of 56) |
| | 6 | 27% (24 of 90) | 19% (7 of 37) |

Data from: A24. n/a: no PV+ neurons were encountered in layer 1 (i.e., denominator = 0).

**Table S4**: GAD1+ and VGAT+ subpopulations of VGLUT1+ neurons, by MERFISH.

| Cell type | Area | N | GAD1+ % (count) | VGAT+ % (count) |
|---|---|---|---|---|
| VGLUT1+ | V1 | 212,063 | 10% (21,755) | 7% (14,888) |
| | V2 | 113,360 | 12% (13,566) | 8% (9,270) |

Data from: A24 (V1 and V2) and A25 (V1 only).

**Table S5**: Co-labeled subpopulations of PV+, SST+, and VIP+ neurons, by 4-FISH.

| Area | N | PV+/SST+/VIP+ % (count) | PV+/SST+ % (count) | PV+/VIP+ % (count) | SST+/VIP+ % (count) |
|---|---|---|---|---|---|
| V1 | 2,595 | 0.08% (2) | 4% (115) | 4% (92) | 0.4% (11) |
| V2 | 959 | 0.4% (4) | 4% (37) | 3% (25) | 1% (10) |

Data from: A24.

**Table S6**: Peptide co-expression by PV+ cells in V1 and V2 by layer, by 4-FISH.

| CBP | Layer | N | % SST+ (count) | % VIP+ (count) |
|---|---|---|---|---|
| PV | 1 | 1 | 0% (0) | 100% (1) |
| | 2/3 | 702 | 6% (45) | 11% (76) |
| | 4/4c | 844 | 7% (60) | 2% (17) |
| | 5 | 77 | 19% (15) | 1% (1) |
| | 6 | 68 | 1% (1) | 0% (0) |

Data from: A24.

**Table S7**: Peptide co-expression by CB+ and CR+ cells in V1 and V2 by layer, by 4-FISH and MERFISH.

| CBP | Layer | % SST+ (count) | % VIP +(count) |
|---|---|---|---|
| CB | 1 | 35% (9 of 26) | 4% (1 of 28) |
| | 2/3 | 53% (4,859 of 9,181) | 3% (340 of 10,215) |
| | 4/4c | 22% (859 of 3,971) | 1% (34 of 4,427) |
| | 5 | 50% (582 of 1,159) | 1% (7 of 1,195) |
| | 6 | 38% (381 of 1,000) | ~0% (4 of 1,021) |
| CR | 1 | 1% (4 of 724) | 7% (52 of 769) |
| | 2/3 | 6% (808 of 12,626) | 32% (4,395 of 13,852) |
| | 4/4c | 22% (873 of 4,021) | 21% (899 of 4,245) |
| | 5 | 30% (284 of 961) | 23% (231 of 998) |
| | 6 | 38% (427 of 1,118) | 17% (192 of 1,147) |

Data from: A24 (V1 and V2), A25 (MERFISH only, V1 only).

**Table S8:** CBP co-expression by peptide+ cells, by layer, 4-FISH and MERFISH.

| Peptide | Layer | % PV+ (count)* | % CB+ (count) | % CR+ (count) |
|---|---|---|---|---|
| SST | 1 | 0% (0 of 1) | 17% (9 of 53) | 8% (4 of 53) |
| | 2/3 | 8% (45 of 540) | 50% (4,859 of 9,780) | 8% (808 of 9,780) |
| | 4/4c | 36% (60 of 169) | 24% (859 of 3,581) | 24% (873 of 3,581) |
| | 5 | 16% (15 of 96) | 22% (582 of 2,654) | 11% (284 of 2,654) |
| | 6 | 1% (1 of 67) | 11% (381 of 3,414) | 13% (427 of 3,414) |
| VIP | 1 | 9% (1 of 11) | 1% (1 of 67) | 78% (52 of 67) |
| | 2/3 | 14% (76 of 539) | 7% (340 of 5,155) | 85% (4,395 of 5,155) |
| | 4/4c | 14% (17 of 125) | 3% (34 of 1,248) | 72% (899 of 1,248) |
| | 5 | 7% (1 of 15) | 2% (7 of 327) | 71% (231 of 327) |
| | 6 | 0% (0 of 3) | 1% (4 of 327) | 59% (192 of 327) |

Data from: A24, A25. * Note that PV data come from A24/4-FISH only.

**Table S9:** Putatively inhibitory subpopulations of SST+, VIP+, CB+, CR+, and VIP+/CR+ neurons, by 4-FISH.

| Cell type | Area | GAD1+ % (count) | VGAT+ % (count) |
|---|---|---|---|
| SST+ | V1 | 80% (309 of 385) | 61% (191 of 312) |
| | V2 | 84% (129 of 153) | 56% (97 of 174) |
| VIP+ | V1 | 62% (411 of 659) | 47% (257 of 552) |
| | V2 | 70% (237 of 338) | 60% (350 of 580) |
| CB+ | V1 | 47% (480 of 1,028) | 45% (330 of 733) |
| | V2 | 50% (239 of 479) | 52% (332 of 635) |
| CR+ | V1 | 51% (467 of 907) | 48% (230 of 477) |
| | V2 | 52% (361 of 695) | 58% (400 of 690) |
| VIP+/CR+ | V1 | 75% (199 of 266) | 56% (95 of 170) |
| | V2 | 81% (145 of 180) | 64% (167 of 261) |

Data from: A24.
